# Thermal adaptation rather than demographic history drives genetic structure inferred by copy number variants in a marine fish

**DOI:** 10.1101/2020.04.05.026443

**Authors:** Hugo Cayuela, Yann Dorant, Claire Mérot, Martin Laporte, Eric Normandeau, Stéphane Gagnon-Harvey, Pascal Sirois, Louis Bernatchez

## Abstract

Increasing evidence shows that structural variants represent an overlooked aspect of genetic variation with consequential evolutionary roles. Among those, copy number variants (CNVs), including duplicated genomic region and transposable elements (TEs) may contribute to local adaptation and/or reproductive isolation among divergent populations. Those mechanisms suppose that CNVs could be important drivers of population genetic structure, whose study is generally restricted to the use of SNPs. Taking advantage of recent developments allowing CNV analysis from RAD-seq data, we investigated how variation in fitness-related traits, local thermal conditions and demographic history are associated with CNVs, and how subsequent copy number variation drives population genetic structure in a marine fish, the capelin (*Mallotus villosus*). We collected 1536 DNA samples from 35 sampling sites in the north Atlantic Ocean and identified 6620 CNVs. We found associations between CNVs and the gonadosomatic index, suggesting that duplicated regions could affect female fitness by modulating oocyte production. We also detected 105 CNV candidates associated with water temperature, among which 20% corresponded to genomic regions located within the sequence of protein-coding genes, suggesting local adaptation to cold water by means of gene amplification. We also identified 175 CNVs associated with the divergence of three parapatric glacial lineages, of which 24% were located within protein-coding genes, which might contribute to genetic incompatibilities and ultimately, reproductive isolation. Lastly, our analyses unveiled a hierarchical, complex CNV population structure determined by temperature and local geography, that was very different from that inferred based on SNPs in a previous study. Our findings underscore the complementarity of those two types of markers in population genomics studies.

## Introduction

Genetic variation is an essential component of evolution, contributing to adaptation, reproductive isolation or genetic structure. Part of this genetic variation, such as single-nucleotide mutation, has been well characterized and its evolutionary role well demonstrated over the last few decades (Morin et al. 2004, Helyar et al. 2011, Leaché et al. 2017). However, there is increasing evidence that structural variants (SVs) represent an overlooked aspect of genetic variation with a possible important evolutionary role (Chain & Feulner 2014, Spielmann et al. 2018, Wellenreuther & Bernatchez 2018, Wellenreuther et al. 2019). A generalized conceptual framework is emerging (Wellenreuther et al. 2019, Mérot et al. 2020), which proposes to study those structural variants and their diversity using integrative approaches to better understand their respective mechanisms and the consequences of their interactions for micro- and macro-evolutionary processes.

Among the different types of SVs, copy number variants (CNVs) have been proposed to play a crucial role in genome evolution, adaptation, and speciation (Freeman et al. 2006, Zhang et al. 2009). CNVs are genomic structural variants in which a segment of DNA can be absent or present in two or more copies due to either gene duplication, deletion, or transposable elements (TEs) (Mérot et al. 2020). CNVs can arise from a variety of mechanisms, including nonallelic homologous recombination, nonhomologous end-joining, and retrotransposition (Hastings et al. 2009). They may be maintained in a population via neutral evolutionary processes (i.e., migration, genetic drift) and either positive, purifying or balancing selection (Zhang et al. 2009, Katju & Bergthorsson 2013, Qian & Zhang 2014).

As copy-number changes may reach high frequency and can be fixed in a short time (Farslow et al. 2015), CNVs have been hypothesized to promote local adaptation and enable the colonization of new habitats (Kondrashov 2012, Qian & Zhang 2014). By affecting gene dosage and expression, gene duplication may determine phenotypic performance and fitness under specific environmental conditions (Kondrashov 2012, Qian & Zhang 2014). For instance, studies reported that gene duplication may underlie adaptive responses to both abiotic (e.g., temperature, heavy metals, and insecticide; Raymond et al. 1991, Hull et al. 2017, Tigano et al. 2018) and biotic factors (e.g., nutriment limitation, Kondrashov et al. 2002). Furthermore, the activity of TEs, a mechanism generating CNVs, can also promote local adaptation (Casacuberta & González 2013, Schrader et al. 2014, Stapley et al. 2015, van’t Hof et al. 2016). TEs generate a great variety of mutations that may lead to phenotypic and fitness variation through the modification of gene expression, the inactivation of genes, and the alteration of gene sequence and reading frame (Chuong et al. 2017). The activation of TEs in response to stress induces structural variation that may help organisms to adapt to environmental conditions (McClintock 1950) such as temperature and rainfall (e.g., Gonzalez et al. 2010).

CNVs may also play a major role in the process of speciation (Lynch & Force 2000, Serrato-Capuchina & Matute 2018). As their evolution rate is higher than that of single nucleotide polymorphisms (SNPs) Sudmant et al. 2013, Paudel et al. 2015, CNVs enhance the accumulation of genetic incompatibilities (Nosil et al. 2009), and thus accelerate reproductive isolation and speciation rate (Böhne et al. 2008, Ricci et al. 2018). TEs may be also involved in reproductive barriers by causing disruptions of gene expression, genomic expansions, and generating new chromosomal inversions (Serrato-Capuchina & Matute 2018).

Clearly, the central role of CNVs in adaptation and reproductive isolation demonstrated in previous studies suggests that structural variants could play a major role in modulating patterns of population structure, whose study is generally restricted to the use of Single Nucleotide Polymorphism (SNPs, Helyar et al. 2011, Hendricks et al. 2018). Given their faster evolution rate (Redon et al. 2006, Sudmant et al. 2013, Paudel et al. 2015) and the effect of the environment on their accumulation within the genome (for TEs, see Chuong et al. 2017), CNVs could reveal patterns of genetic structure that are different from those drawn by SNPs. Consequently, they would provide a valuable and complementary set of genetic markers to analyze neutral and adaptive population structure in both basic and applied studies. However, with the exception of targeted candidate genes studies, CNV genotyping remained expensive for a long time as it required whole genome sequencing data with deep coverage (Pirooznia et al. 2015, Makałowski et al. 2019), precluding the use of these markers in population genomic studies that involve thousands of samples from dozens of populations. This situation has recently changed through the development of a novel methodological approach allowing the identification of CNVs in a cost-effective way using RAD-seq data (McKinney et al. 2017, Dorant et al. 2020). Using RAD-seq data, Dorant et al. (2020) recently showed that CNVs can be more efficient than SNPs at revealing genotype-temperature associations in marine invertebrates. Yet, at present time, the usefulness of CNVs from RAD-seq data as valuable DNA markers for population genomic studies still needs to be more broadly exemplified.

The goal of this study was to investigate how fitness, local thermal conditions and demographic history are associated with CNVs genotyped in multiple populations, and subsequently how copy number variation drives population genetic structure in a marine fish, the capelin (*Mallotus villosus*), an excellent biological model to address this issue. First, capelin has a complex demographic history and in the north Atlantic it comprises three glacial lineages (NWA, ARC, and GRE) that diverged in allopatry from 1.8 to 3.5 MyA (Cayuela et al. 2019). Very low migration rates and the absence of admixture between lineages despite no obvious physical barriers suggest the existence of strong reproductive barriers and an ongoing speciation process (Cayuela et al. 2019), allowing the consideration of copy-number variation at two taxonomic scales. Second, within-lineage populations may reproduce in either demersal or beach-spawning sites (Christiansen et al. 2008). In the latter case, populations span large environmental gradients in which temperature seems to be an important selective driver affecting fitness components (i.e., survival and growth rate) at early stages (i.e., embryo and larvae; Frank & Leggett 1981, Leggett et al. 1984) and probably later in life (Colbeck et al. 2011, Kenchington et al. 2015). In the NWA lineage, genotype-temperature associations based on SNPs suggest that beach-spawning individuals are locally adapted to thermal conditions prevailing in the intertidal zone (Cayuela et al. 2019).

More precisely, we first used the method developed by McKinney et al. (2017) and refined by Dorant et al. (2020) to detect reliable CNVs. Second, we investigated how CNVs normalized read depth, a robust proxy of putative copy number (Dorant et al. 2020), correlates with the gonadosomatic index, a commonly used fitness proxy in fishes (Brewer et al. 2008, Ressel et al. 2019). Third, we examined the putative role of CNVs in thermal adaptation by assessing correlations between normalized read depth and water temperature in beach spawning sites. Based on previous findings in Antarctic notothenioid fishes (Chen et al. 2008), we expected a higher number of gene copies in individuals from cold waters than in their counterparts from warmer waters. According to recent findings in teleost fishes (Yuan et al. 2018, Carducci et al. 2019, Shao et al. 2019), we also hypothesized that thermal shocks could have led to temperature-dependent accumulation of TEs in the capelin genome. Fourth, we investigated the potential role of CNVs on the ongoing speciation process among glacial lineages by quantifying how the normalized read depth differs for each pair of lineages. We expected that variation in copy number of protein-coding genes and the rapid accumulation of class I (i.e., retrotransposons) and class II (i.e., DNA transposons) TEs could be involved in the reproductive isolation responsible for the absence of admixture between lineages (Cayuela et al. 2019). Fifth, we examined how variation in copy number associated with thermal conditions and demographic history drive genetic structure in the study area. Lastly, we underscored the differences in the patterns of genetic structure drawn by CNVs and SNPs, discussed the evolutionary mechanisms driving those discrepancies, and emphasized the complementarity of both markers in the context of population genomic studies.

## Materials and methods

### Sampling area, phenotypic analyses

We sampled 1536 capelins from 35 spawning sites in the Northwest Atlantic, in Canadian and Greenland waters (**Fig.1** and **Supplementary material S1**, **Table S1**). Three sites were sampled in the ARC lineage, four sites in GRE lineage, and 28 sites in the NWA lineage. In this last lineage, the sites included 18 beach spawning sites, six demersal spawning sites, and four offshore sites. In the whole dataset, the median sample size was 46.5 (range: 19 to 50) individuals per site. The fish were collected and immediately frozen, and a piece of fin was preserved in RNAlater. The gonadosomatic index was measured at the laboratory on 843 individuals (302 females and 541 males) from the 18 beach spawning sites of the NWA lineage.

**Fig.1.**
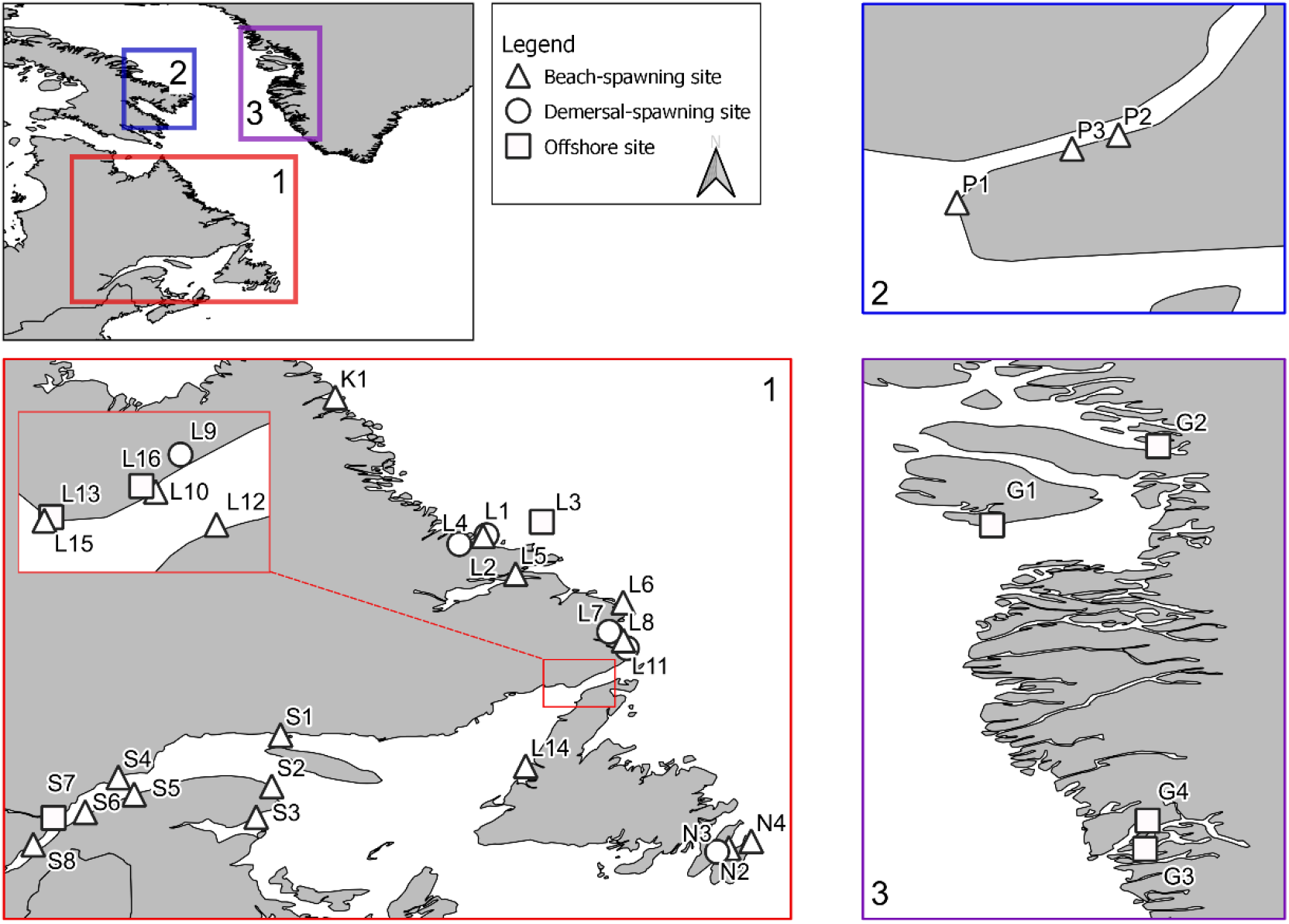
Map of the study area showing the sites sampled in the three glacial lineages. (1) Sampling sites in the NWA lineage. (2) Sampling sites in the ARC lineage. (3) Sampling sites in the GRE lineage. Three types of sites were sampled: beach-spawning sites (triangle), demersal-spawning sites (circle), and offshore sites (square).

### DNA sequencing, genotyping, and CNV discovery

Both DNA extractions and library preparations were performed following protocols fully described in Cayuela et al. (2020). Libraries were size-selected using a BluePippin prep (Sage Science), amplified by PCR and sequenced on the Ion Proton P1v2 chip (single-end sequencing). Eighty-two individuals were sequenced per chip.

Barcodes were removed using cutadapt (Martin 2011) and trimmed to 80 bp, allowing for an error rate of 0.2. They were then demultiplexed using the ‘process_radtags’ module of Stacks v1.48 (Catchen et al. 2013) and aligned to the capelin draft genome (Cayuela et al. 2019) assembly using bwa-mem (Li 2013) with default parameters, as detailed below. Next, aligned reads were processed with Stacks v.1.48 for SNP calling and genotyping. The ‘pstacks’ module was used with a minimum depth of 3 and up to 3 mismatches were allowed in the catalog creation. We then ran the ‘populations’ module to produce a vcf file that was further filtered using python (https://github.com/enormandeau/stacks_workflow) and bash scripts. SNPs were kept if they displayed a read depth greater than 4 and less than 70. Then, SNPs present in at least 70 % of individuals in each sampling location and with heterozygosity lower than 0.60 were kept to further control for paralogs.

The *HDplot* approach proposed by McKinney et al. (2017) allows identifying CNVs based on two characteristics: the proportion of heterozygous individuals within a population and allelic ratios within heterozygous individuals. We used a refined version of this approach (Dorant et al. 2020) that discriminates “singleton” SNPs (i.e., non-duplicated) from “duplicated” SNPs (i.e., combining duplicated SNPs and SNPs with a high coverage) using 5 parameters: (1) proportion of heterozygotes (PropHet), (2) Fis, (3) median of allele ratio for heterozygotes (MedRatio), (4) median of SNP read depth for heterozygotes (MedDepthHet) and (5) median of SNP read depth for homozygotes (MedDepthHom). The parameters were calculated using a custom python script (available at https://github.com/enormandeau/stacks_workflow) parsing the filtered VCF file. The five parameters were plotted pairwise to visualize their distribution across all loci. Based on the graphical demonstration by McKinney et al (2017) and the approach proposed by Dorant et al. (2020), we considered different combinations of parameters and graphically fixed the cut-off of the four categories of SNPs (singleton SNPs, duplicated SNPs, high coverage SNPs, and low confidence). We then kept one single marker per locus (the one with the highest minor allele frequency) and extracted the read depth of duplicated loci to construct the CNV dataset using vcftools. Therefore, for more simplicity, the term “CNV” is used to define all loci classified as “duplicated” and “high coverage” by our approach. Following the procedure described in Dorant et al. (2020), CNV locus read counts were normalized to account for differences in sequencing effort across all samples. Normalization was performed using the Trimmed mean of M-values method originally described for RNA-seq count normalization and implemented in the R package edgeR (Robinson & Oshlack 2010). The correction accounts for the fact that for an individual with a higher copy number at a given locus, that locus will contribute proportionally more to the sequencing library than it will for an individual with lower copy number at that locus. This procedure was applied to the whole dataset (i.e., the 1536 capelins from 35 spawning sites) to detect the CNVs present throughout the study area.

Using the capelin reference genome (Cayuela et al. 2019), we examined if the CNVs discovered were located within sequences of protein genes or within intergenic regions. Note that the genome is assembled at the scaffold level; nevertheless, contig order was reconstructed via synteny analysis, which allowed the sorting of contigs into 24 orthologous chromosomes (Cayuela et al. 2019). For CNV candidates located in intergenic regions, we examined whether they corresponded to repeated elements including TEs. For that purpose, the reference genome was annotated for TEs (both DNA transposons and retrotransposons) and interspersed repeats (including satellites, simple repeats, and low complexity DNA sequences) using RepeatMasker 4.1 (Smit et al. 2015) with the default options and the dfam database (Hubley et al. 2016) for the zebrafish (*Danio rerio*).

### Investigating CNV-fitness associations

To investigate the potential effect of CNVs on fitness related traits, we examined how gonadosomatic index correlates with normalized CNV read depth using a locus-by-locus GWAS-like approach. The gonadosomatic index, expressed as gonad mass as a percentage of total body mass, is widely used as a simple measure of the extent of reproductive investment and gonadal development (Gunderson 1997, Brewer et al. 2008, Ressel et al. 2019). As the gonad mass strongly differs between sexes, we analyzed each sex (302 females and 541 males) separately. We used Linear Mixed Models (LMMs) with restricted maximum likelihood optimization where the log-transformed gonadosomatic index was introduced as the response variable and the scaled normalized read depth was included as the explanatory variable. The spawning site was included as a random effect (i.e., random intercepts) to account for the non-independency of observations across sites. One model was performed for each CNV locus. We used likelihood ratio tests (comparing the models with and without the explanatory term) to assess the significance of the CNV-fitness associations. The analysis was conducted in the R package lme4 (Bates et al. 2015). We used a false discovery rate (FDR) of 0.10 to limit the risk of type I error when conducting multiple comparisons.

### Investigating CNV-temperature and CNV-lineage associations

To investigate the potential role of CNVs in local adaptation, we examined whether normalized CNV read depth (i.e., a robust proxy of putative copy number, Dorant et al. 2020) correlates with sea water temperature, an environmental factor affecting growth and survival of embryos and larvae (Frank & Leggett 1981, Leggett et al. 1984). The marine data layer for bottom annual temperature was downloaded from Bio-ORACLE (http://www.bio-oracle.org/) and temperature at spawning sites was extracted using the R package sdmpredictors (Bosch et al. 2017). CNV-temperature associations were evaluated by combining locus-by-locus regressions and multivariate analyses. We ran LMMs with restricted maximum likelihood optimization where the log-transformed normalized read depth was incorporated as the response variable and temperature as the explanatory variable. The spawning site was included as a random effect (i.e., random intercepts) to account for the non-independency of observations across sites. One model was performed for each CNV, and we used likelihood ratio tests and a FDR of 0.10 following the method of Benjamini & Hochberg (1995) to assess the significance of CNV-temperature associations.

We also used a partial redundancy analysis (pRDA, Legendre & Legendre 2012) to detect multi-CNV candidates that were associated with temperature after controlling for the spawning sites, as commonly done for SNP data (Laporte et al. 2016, Le Luyer et al. 2017, Forester et al. 2018 for details). Global and marginal analyses of variance (ANOVA) with 1,000 permutations were performed to assess the significance of the models and evaluate the contribution of each environmental variable. Once CNVs were loaded against the pRDA axes, candidates for an association with temperature were determined as those exhibiting a loading greater than 2.25 standard deviations from the mean loading (P < 0.01) (Forester et al. 2018). We retained the outliers that were detected by both LMMs and pRDA to reduce the number of false positives and represented the overlap with a Venn diagram.

We used a similar approach to explore the potential role of CNVs in the ongoing speciation process among the three aforementioned glacial lineages (Dodson et al. 2007, Cayuela et al. 2019). The three lineages (ARC, GRE, and NWA) diverged approximately from 1.8 MyA (NWA-GRE) to 3.8 MyA (ARC-NWA), display limited historical introgression, and exhibit no apparent contemporary admixture (Cayuela et al. 2019). The procedure was performed separately for each pair of lineages. In LMMs, the lineage was introduced as a discrete explanatory variable with two modalities.

We then examined whether the CNV candidates were located within sequences of protein-coding genes or within intergenic regions and if so, whether they corresponded to TE (i.e., DNA transposons and retrotransposons) or interspersed repeats (including satellites, simple repeats, and low complexity DNA sequences).

### Inferring genetic structure based on CNVs

We documented population structure based on CNVs using a hierarchical clustering analysis (Langfelder & Horvath 2012). We first calculated Bray-Curtis distance for each pair of individuals as commonly used in landscape genetic studies (Shirk et al. 2014). Then, we calculated the mean of the individual genetic distances for each pair of sites to obtain a square matrix of between-site genetic distances. With this distance matrix, we performed hierarchical clustering analyses using the R function *hclust*, based on the Ward’s minimum variance method (option *ward.D2*, Murtagh & Legendre 2014). To evaluate the robustness of the dendrogram clusters, we performed 10,000 bootstraps using the R function *boot.phylo* implemented in the *ape 5.3* package (Paradis et al. 2019). Dendrogram nodes were considered robust if their bootstrapping value was > 0.80. Then, we examined how temperature of beach-spawning sites may affect the genetic structure inferred from CNVs within the NWA lineage. We performed two hierarchical clustering analyses: one based on all 6620 CNVs detected and the other with the 105 CNV candidates associated with temperature (see Results). Then, we examined how lineage demographic divergence affects the genetic structure inferred from CNVs. To maintain a relatively balanced number of sites in the three lineages (NWA, GRE, and ARC), we selected six spawning sites within the NWA lineage based on the clustering analysis performed for this lineage: three sites represented genetic cluster 1 (L2, L6, and L10) and three sites represented cluster 2 (S1, S2, and S3) (see Results). Two hierarchical clustering analyses were conducted: one based on all 6620 CNVs and the other with the 175 CNV candidates associated with lineage divergence (see Results).

The clustering analysis for the 18 beach-spawning sites within the NWA lineage revealed two genetic clusters: cold sites from Labrador and the Atlantic coast of Newfoundland were grouped in cluster 1 whereas warm sites from the Gulf/estuary of St. Lawrence R. were grouped in cluster 2 (see Result section). Interestingly, the spawning site S8 located in the St. Lawrence R. estuary was assigned to cluster 1. We thus hypothesized that if the individuals of site S8 are adapted to the cold waters of the Saguenay fjord (see Discussion section), they should have higher normalized read depth than individuals from the warm waters of the nearby Gulf of St. Lawrence. We tested this hypothesis by building linear models where the log-transformed normalized read depth was included as the response variable and the type of site (S8 versus the geographically close sites from the Gulf of St. Lawrence S1, S2, S3, S4, S5, and S6) as the explanatory variable. One model was performed for each CNV, and we used likelihood ratio tests and a stringent FDR (0.001) following the method of Benjamini & Hochberg (1995) to assess the significance of CNV-temperature associations.

## Results

### Sequencing statistics and CNV discovery

For the whole dataset (1536 capelins from 35 spawning sites), GBS produced 897,382 ± 461,911 reads per sample on average before any quality filtering. The SNP calling process identified 642,098 SNPs that were successfully genotyped in at least 70% of the samples, with a low rate of missing data (median of 1.8%). The *HDplot* approach identified 280 duplicated markers and 14,319 markers with high coverage distributed over 6620 loci (hereafter called CNVs). Moreover, 48,591 markers were classified as low confidence, and 578,819 markers were classified as singleton SNPs (without any filtering).

Among the 6620 CNV loci, 1519 (22%) were located within protein-coding genes and 327 (5%) corresponded to repeated elements (**Fig.3A**). Among the latter, 50% were retrotransposons (i.e., LRT and non-LRT retrotransposons; for the specific families of retrotransposons, see **Supplementary material S1**, **Table S2**). Moreover, 31% of the repeated elements corresponded to interspersed elements including simple repeats, low complexity regions, and satellites (**Supplementary material S1**, **Table S3**). The other repeated elements were DNA transposons (14%) and small RNAs (5%) (**Supplementary material S1**, **Table S3**).

### Associations between CNVs and fitness proxy

We detected six CNV candidates (**Table 1**) associated with the gonadosomatic index in females whereas no strong candidates were detected for males (**Supplementary material S1**, **Fig.S1**). In females, the CNV candidates were located on chromosomes 6, 9, 11, 16, and 19 (**Fig.2**). The candidate 50008_8 was found within the sequence of a gene (**Table 1**) with unknown function. The CNV 48830_43 was included within a 5-Kbp window around a gene involved in the regulation of various cellular processes (**Table 1**). The candidate 25048_60 was in an intergenic region at a distance of ~20 kbp from a gene (XM_012823774.1) that regulates the activation of the follicle-stimulating hormone (FSH) receptor (**Table 1**), which directly controls female oocyte production in vertebrates. The relationship between the gonadosomatic index and the normalized read depth of this candidate was negative (slope coefficient: −0.15±0.03, *R*^2^ = 0.11), suggesting that a high number of copies of the gene (or its promotor) regulating the receptor of the FSH could decrease female fecundity.

**Table 1.** Candidate CNVs associated with gonadosomatic index. We provide the CNV name (CNV ID), the contig name and gene name from the Capelin genome assembly, the chromosome name inferred using synteny analyses by Cayuela et al. (2020; U = unanchored contig), the type of repeated elements, and the gene Pfam. We retained protein-coding genes if the candidate CNV was inside the gene sequence (i.e., XM_012828078.1) or within a 5-Kbp window around the gene (XM_012836008.1). We also present the gene XM_012823774.1 (distance of ~20Kbp from the candidate 25048_60) because of its potential direct effect on the gonadosomatic index.

| CNV ID | Contig | Chr | DNA repeated elements | Gene name | Gene Pfam |
| --- | --- | --- | --- | --- | --- |
| 50008_8 | contig_5 | 6 | None | XM_012828078.1 | None |
| 59193_13 | contig_88 | 16 | None | None | None |
| 25048_60 |  | 11 | None | XM_012823774.1 | PF00001; PF12369; PF13306; |
|  | contig_306 |  |  |  | PF13855 |
| 48830_43 | contig_5712 | 19 | None | XM_012836008.1 | PF04752 |
| 53320_53 | contig_69 | U | None | None | None |
| 31892_56 | contig_358 | 4 | None | None | None |

**Fig.2.**
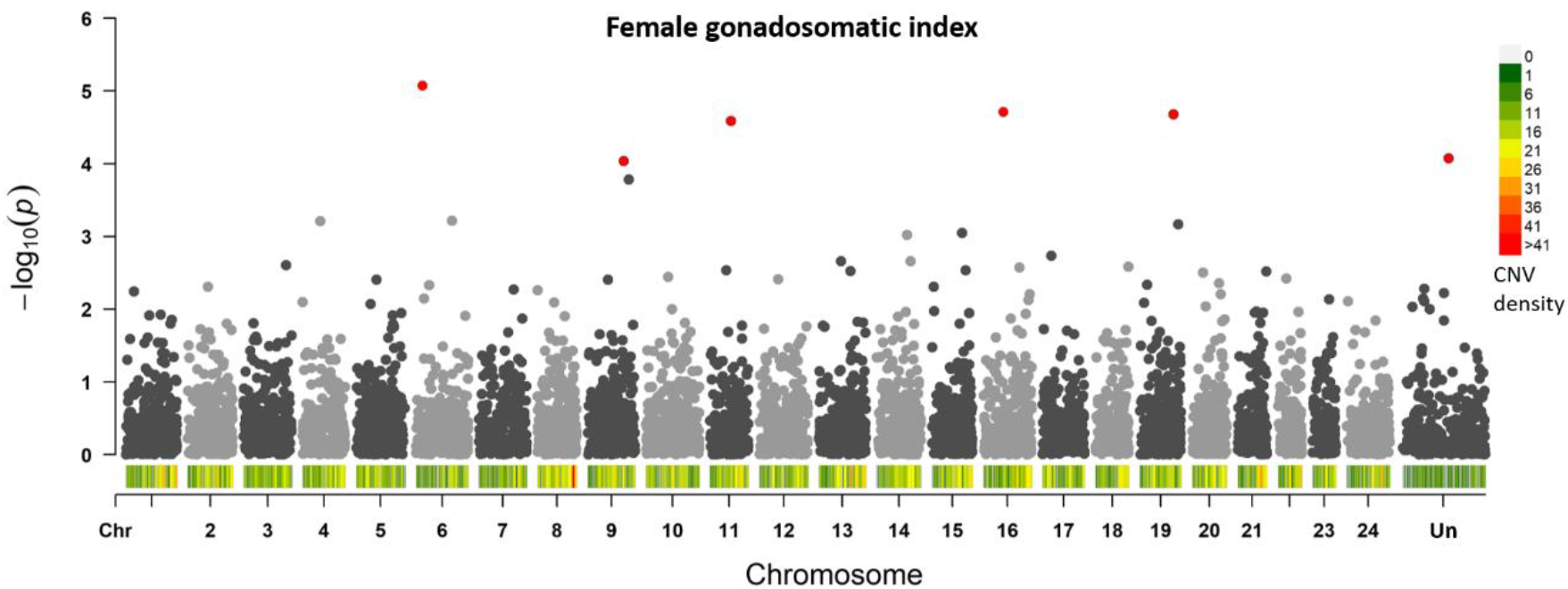
Manhattan plot showing the distribution of candidate CNVs associated with female gonadosomatic index. The p-values (-log(p-value) of LRT tests associated with LMMs are shown on the Manhattan plot. Red dots indicate the candidate CNVs determined using both pRDA and LMMs

**Fig.3.**
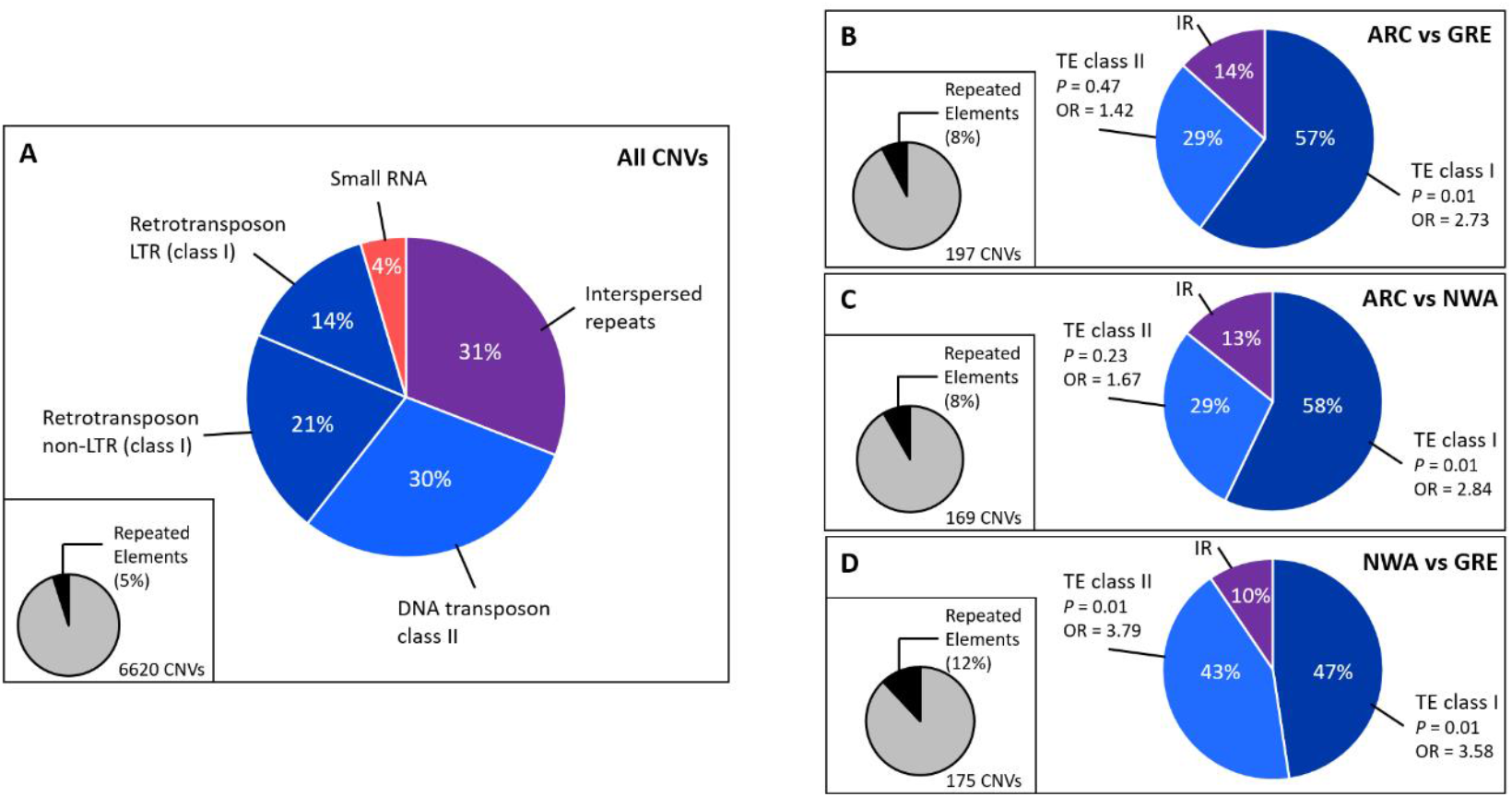
Repeated elements among the 6620 CNVs detected in the entire data set. (A) Type of repeated elements (i.e., interspersed repeats, DNA transposons, retrotransposons LTR and non-LTR, and small RNAs). Interspersed repeats include simple repeats, DNA satellites, and low complexity regions. (B-D) TEs in the set of candidate CNVs detected for each pair of lineages (ARC-GRE, ARC-NWA, and NWA-GRE). The proportion of repeated elements among candidate CNVs is also presented, as well as the composition of three classes of repeated elements: TE class I (i.e. retrotransposons, LTR and non-LTR), TE class II (i.e., DNA transposons), and interspersed repeats (IR). The results of the Fisher tests performed to examine an excess of TE classes I and II in candidate CNVs are provided (*p*-value and odd ratio, OR).

### Associations between CNVs and temperature

The pRDA (**Fig.4A**) built to detect CNVs associated with temperature was highly significant (*df* = 1, *F*-statistic = 26.79, *p* = 0.001), although the coefficient of determination was low (adjusted *r*^2^ = 0.03). The pRDA detected 106 candidate CNVs associated with temperature, whereas LMMs identified 1932 candidate CNVs (**Fig.4B**). A total of 105 CNVs related to temperature were common to the two methods and thus considered as strong candidates (**Fig.4B**).

**Fig.4.**
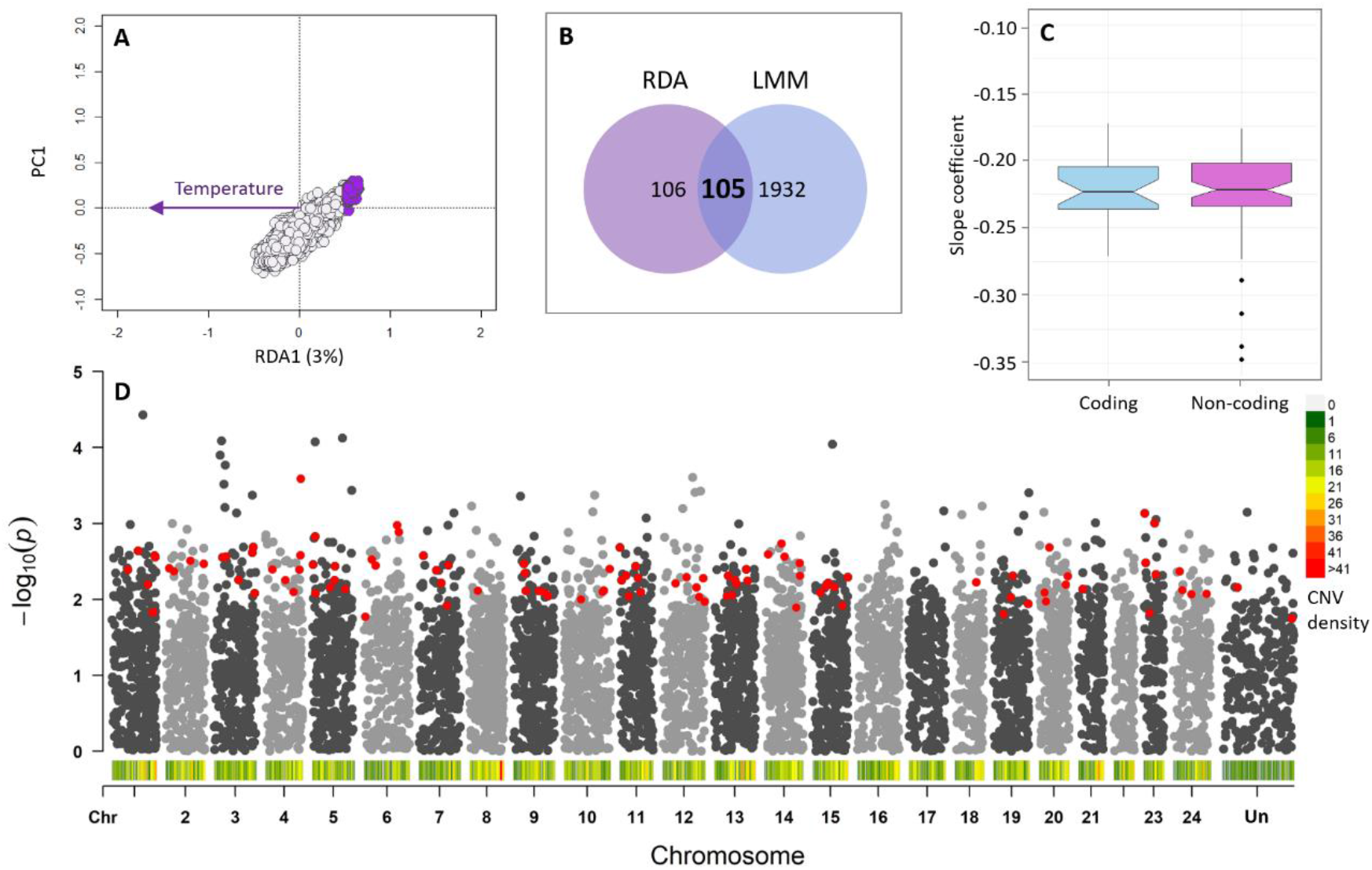
CNV-temperature associations in the Capelin. (A) pRDA including temperature: purple dots show candidate CNVs for temperature. (B) The Venn diagram shows the number of candidate CNVs associated with temperature detected using the pRDA, LMMs, and both methods combined (intercept). (C) Slope coefficients from LMMs for the effect of temperature in the 105 candidate CNVs: coefficients are always negative for both coding and non-coding regions (for regression outputs, see **Supplementary material S1**, **Table S3)**. (D) Manhattan plot showing the distribution of candidate CNVs associated with temperature along the Capelin genome. The p-values (-log(p-value) of LRT tests associated with LMMs are shown on the Manhattan plot. Red dots indicate the candidate CNVs detected by both pRDA and LMMs.

Those CNVs were spread among all chromosomes except 16, 17, and 22 (**Fig.4D**). For all 105 candidate CNVs, correlations between normalized read depth and temperature were negative (for the slope coefficients, see **Supplementary material S1**, **Table S3**), indicating that the number of copies decreased with temperature in both coding and non-coding regions (**Fig.4C**). Twenty-eight candidate CNVs (23%) were located within the sequence of protein-coding genes (see **Table S4)**. Fisher tests showed that those candidates were not present in excess within protein-coding genes (odd ratio = 1.12, *p* = 0.75). Furthermore, our analyses revealed that six of those candidate CNVs (6%) corresponded to repeated elements. Four of them were DNA transposons, one was a retrotransposon (LTR, Gypsy family), and one corresponded to simple repeats. The low number of repeated elements precluded rigorous statistical analysis of their composition or their potential excess in representation.

### Associations between CNVs and glacial lineages

The pRDA performed to identify CNVs associated with glacial lineages was highly significant for the pairs of lineages ARC-GRE (*df* = 1, *F*-statistic = 10.05, *p* = 0.001), ARC-NWA (*df* = 1, *F*-statistic = 17.03, *p* = 0.001), and NWA-GRE (*df* = 1, *F*-statistic = 18.01, *p* = 0.001). We detected 205 candidate CNVs associated with the pair of lineages ARC-GRE, 170 candidates for ARC-NWA, and 183 for NWA-GRE (**Fig.5**). Yet, the coefficient of determination was relatively low in the three cases (ARC-GRE: adjusted *r*^2^ = 0.03; ARC-NWA: adjusted *r*^2^ = 0.01; and NWA-GRE: adjusted *r*^2^ = 0.01).

**Fig.5.**
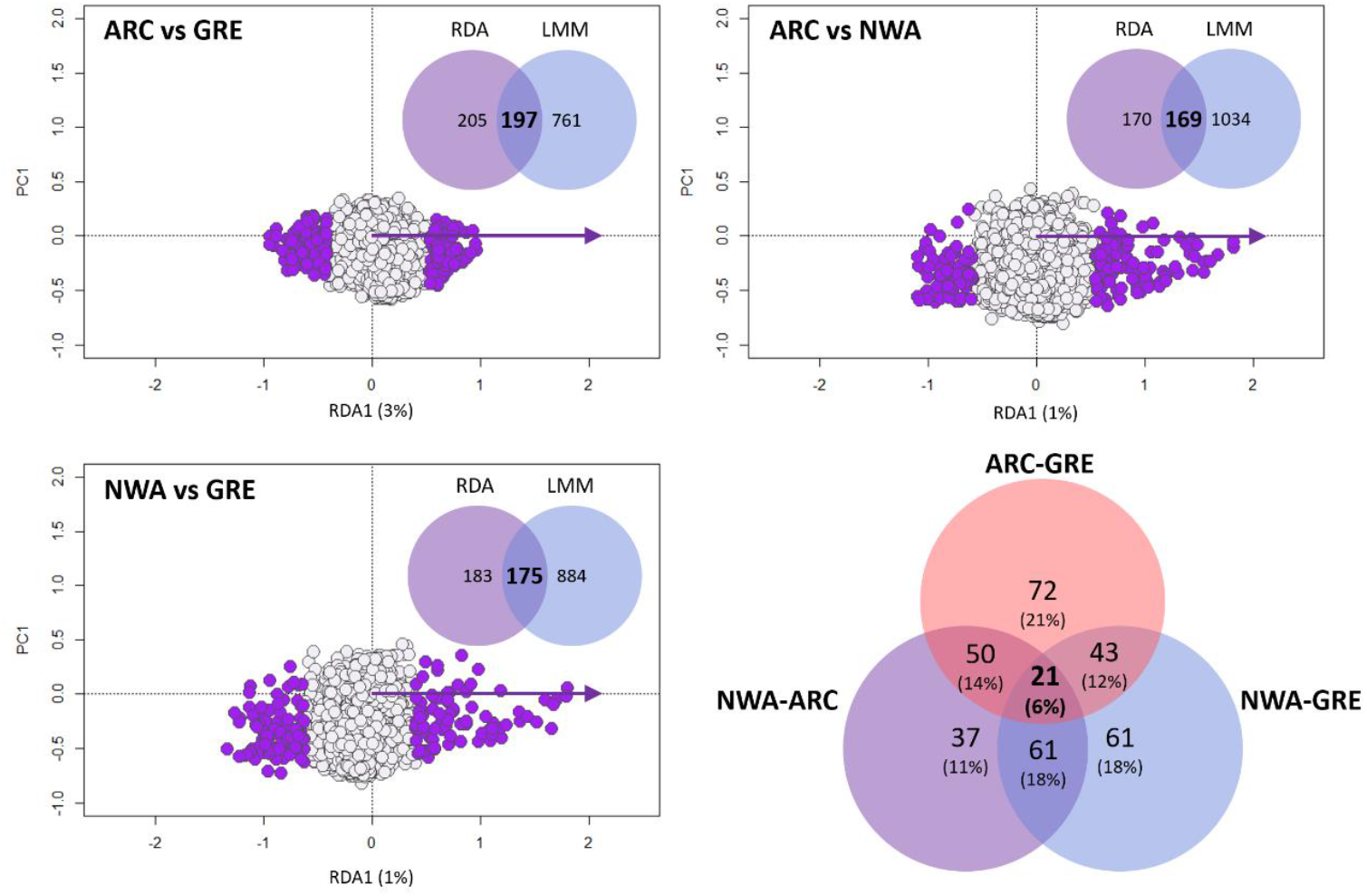
CNV-lineage associations in the Capelin. pRDAs include the lineage as explicative variable: purple dots show candidate CNVs. Venn diagrams for each of the pRDAs show the number of candidate CNVs associated with lineages detected using the pRDA, LMMs, and both methods combined. The Venn diagram on the right shows the number of candidate CNVs shared between or private to each pair of lineages

Using LMMs, we identified 761 candidate CNVs associated with the divergence between lineages ARC and GRE, 1034 candidates between ARC and NWA, and 884 between NWA and GRE (**Fig.5**). Among those, 197 were detected by both pRDA and LMMs for the pair of lineages ARC-GRE, 169 for the pair ARC and NWA, and 175 for the pair NWA and GRE, making them strong candidates associated with divergence between these lineages (**Fig.5**). Twenty-one of those CNVs were common to all three pairwise comparisons (**Fig.5**). Candidate CNVs were spread across the 24 chromosomes regardless the pair of lineages considered (**Fig.6**).

**Fig.6.**
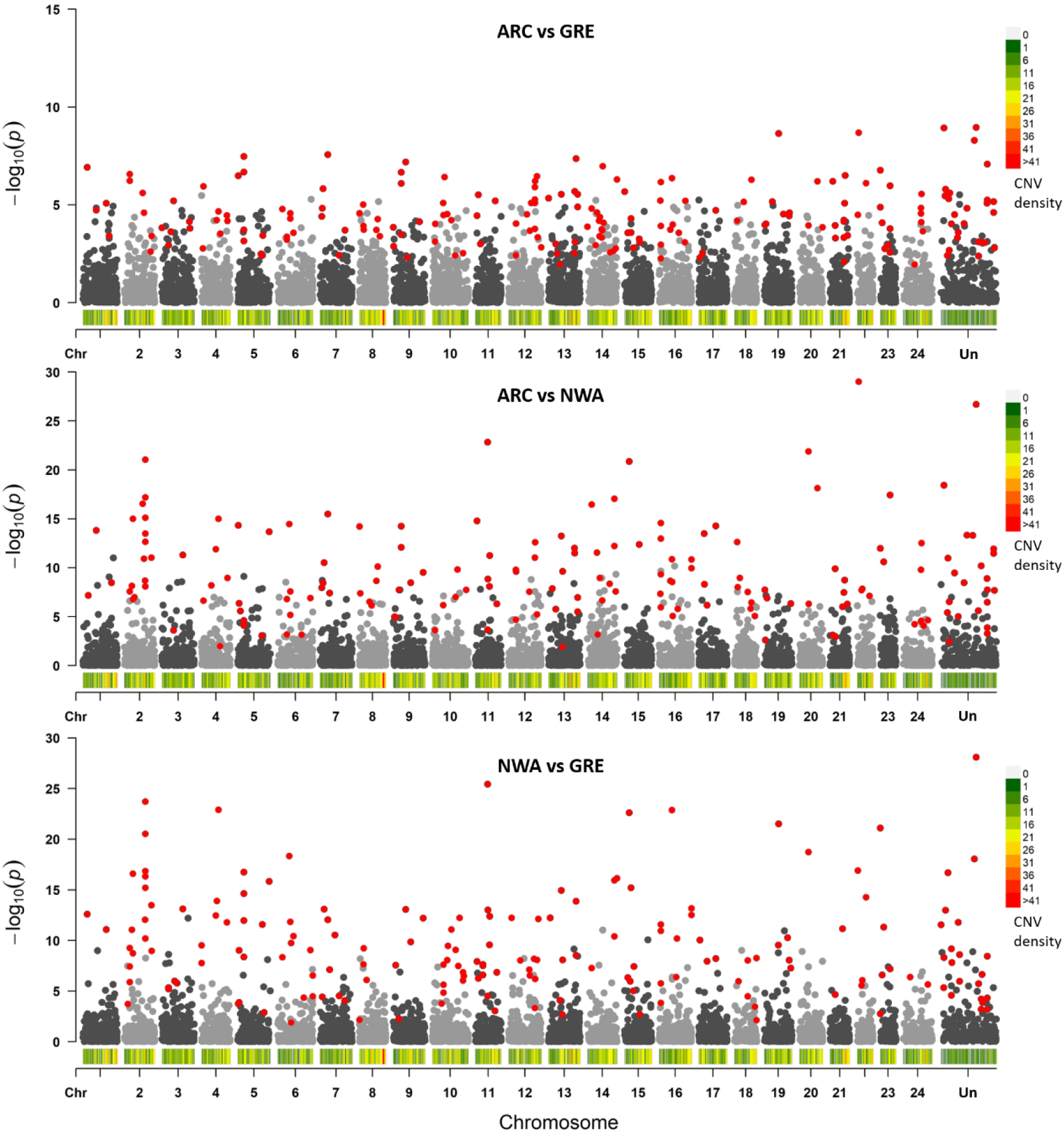
Manhattan plot showing the distribution of candidate CNVs associated with the three pairs of Capelin glacial lineages (i.e., ARC-GRE, ARC-NWA, and NWA-GRE). The p-values (-log(p-value) of LRT tests associated with LMMs are shown on the Manhattan plot. Red points indicate the candidate CNVs determined using both pRDA and LMMs.

Forty-six (23%), 43 (25%), and 30 (17%) candidate CNVs were located within protein-coding genes in the pairs of lineages ARC-GRE, ARC-NWA, and NWA-GRE respectively (see **Table S5**). Fisher tests indicated that these candidates were not present in excess in the sequences of protein-coding genes (ARC-GRE: odd ratio = 0.75, *p* = 0.94, ARC-NWA: odd ratio = 1.11, *p* = 0.30, NWA-GRE: odd ratio = 1.02, *p* = 0.48).

Eight percent of candidate CNVs were repeated elements for the ARC-GRE and ARC-NWA lineage pairs, and 12% for NWA-GRE (**Fig.1B-D**; see the detailed composition of those repeated elements in **Supplementary material S1**, **Table S6**). We also found that the repeated elements were mainly DNA transposons and retrotransposons (**Fig.5**). Fisher tests revealed that the candidate CNVs of the three lineage pairs were enriched for class I TEs (**Fig.5**). That is, both LRT and non-LRT retrotransposons were ~3 times more abundant in the set of candidate CNVs than in entire set of 6620 CNVs. By contrast, DNA transposons were found in excess (~4 times higher than expected) for the NWA-GRE lineage pair only (**Fig.5**).

### Hierarchical genetic structure based on CNVs

Our findings suggest that the significant association between CNVs and temperature could reflect adaptive genetic structure within the NWA lineage. Hierarchical clustering analyses (based on the 6620 CNVs) revealed the existence of two genetic clusters, one occurring in the northern part of study area (cluster1) along the Atlantic coast and the other occupying the Gulf of St. Lawrence (cluster 2) (**Fig.7A** and **Fig.8**). The CNV divergence between the two clusters was strongly supported by the data as reflected by the 100% bootstrap support for the first node of the dendrogram (**Fig.7A**). Cluster 1 (9 spawning sites) occurs in relatively cold waters, with a mean bottom temperature of 1.71±1.33°C (min: −0.13, max: 2.82). By contrast, cluster 2 (9 spawning sites) occurs in warmer waters with a mean bottom temperature of 4.23±1.47°C (min: 2.09, max: 5.97). The high bootstrap support (> 80%) at the shallower nodes of the dendrogram (**Fig.7A**) suggested a complex genetic substructure within the two clusters associated with geographical proximity and/or thermal similarity of spawning sites. Note that the hierarchical clustering analyses based on the whole set of CNVs (*n* = 6620) or the 105 candidates associated with temperature were very similar, suggesting that temperature-dependent CNVs drive the genetic structure within the NWA lineage.

**Fig.7.**
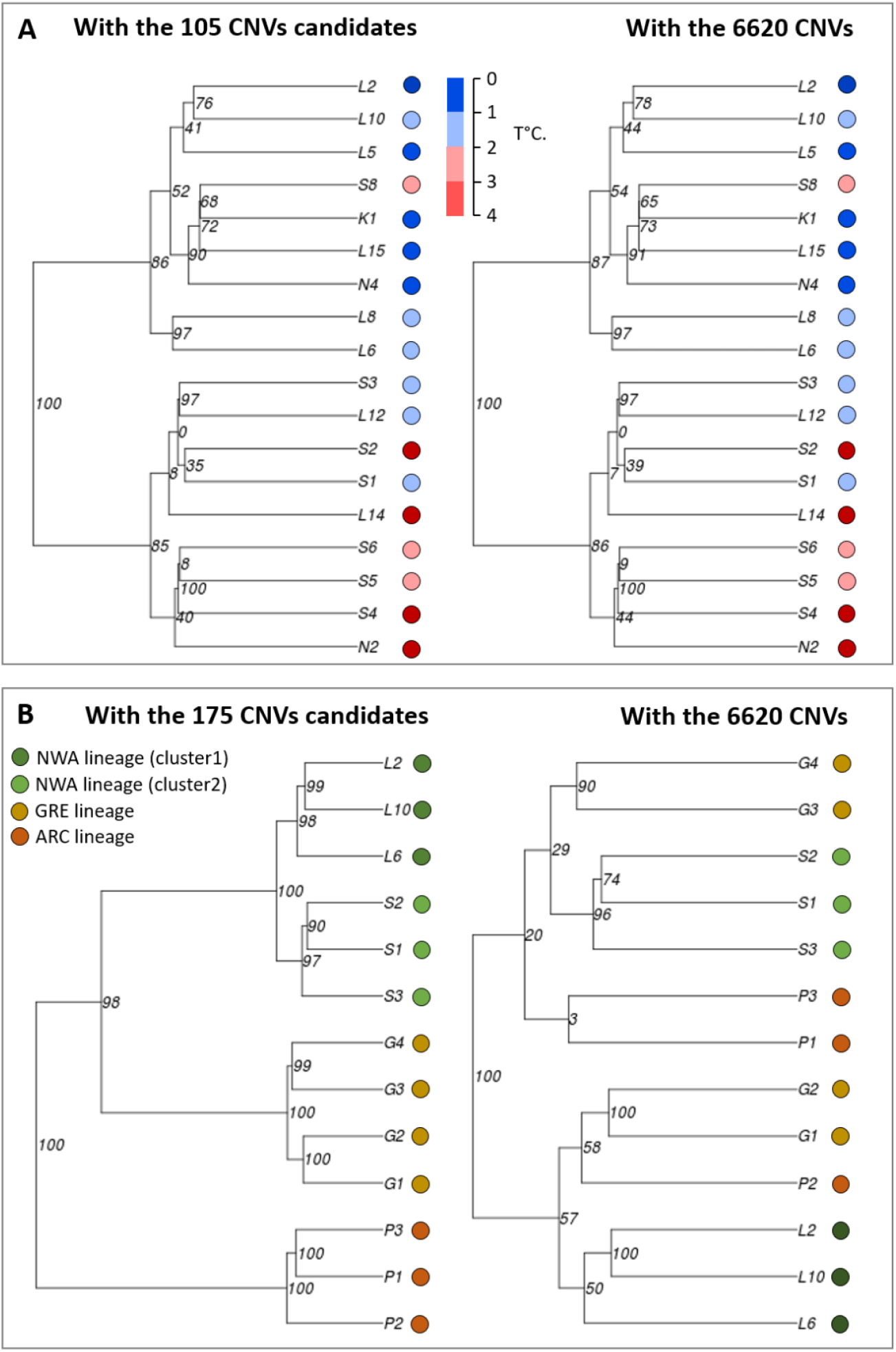
Dendrograms from hierarchical clustering analysis based on the Ward’s minimum variance method revealing genetic structure based on CNVs. To evaluate the robustness of the dendrogram clusters, we performed 10,000 bootstraps and dendrogram nodes were considered to be robust if their bootstrapping value was > 0.80. (A) We investigated how temperature of beach-spawning sites may affect the genetic structure inferred from CNVs within the NWA lineage. We performed two hierarchical clustering analyses: one based on the 6620 CNVs and the other with the 105 candidate CNVs associated with temperature. (B) We examined how lineage demographic divergence affects the genetic structure inferred from CNVs. To have a relatively balanced number of sites in the three lineages, we selected six spawning sites within the NWA lineage based on the clustering analysis performed for this lineage (A): three sites represented the genetic cluster 1 (L2, L6, and L10) and three sites the cluster 2 (S1, S2, and S3). Two hierarchical clustering analyses were conducted: one based on the 6620 CNVs and the other with the 175 candidate CNVs associated with lineage divergence. See Fig. 1 for sampling locations.

**Fig.8.**
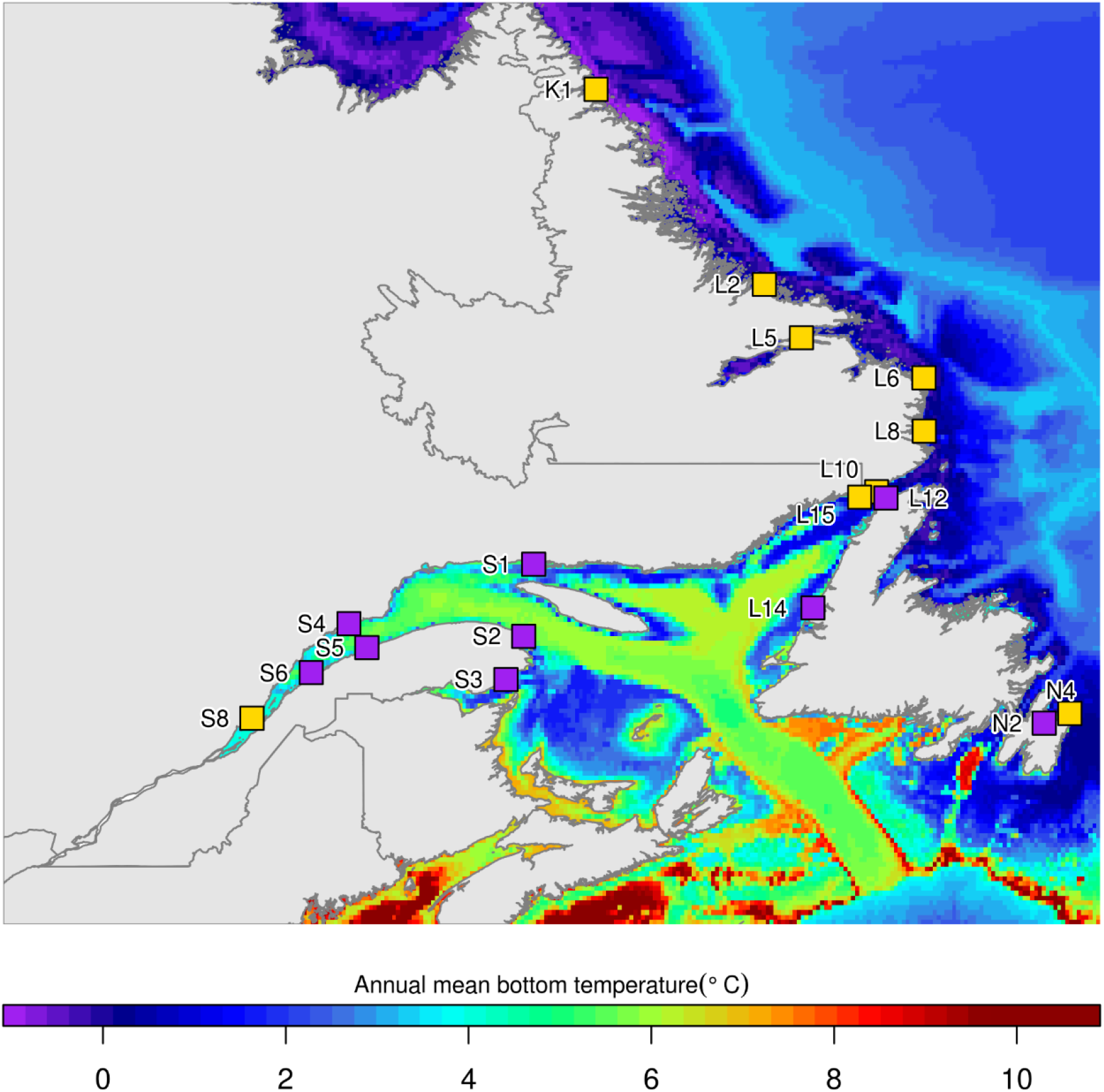
Map showing the genetic structure inferred by CNVs for the 18 beach-spawning sites of the NWA lineage. Yellow and purple squares show the spawning sites grouped within genetic clusters 1 and 2 respectively.

Regardless the set of the CNVs used, the clustering analysis indicated that the site S8 in the St-Lawrence R. estuary was assigned to cluster 1 despite its closer geographic proximity with other sites within cluster 2 (**Fig.8**). Regression analyses showed that capelins from this site have a higher normalized read depth than those from spatially close sites of the Gulf of St. Lawrence R. for 104 of the 105 candidate CNVs associated with temperature (**Supplementary material S1**, **Table S7**). This pattern supports the hypothesis that individuals from the Saguenay fjord are more genetically related to the cold-adapted capelins from cluster 1 than the warm-adapted capelins from the Gulf of St. Lawrence.

Lastly, we investigated whether the demographic divergence of glacial lineages based on CNVs contributed to the genetic structure across the study area. Hierarchical clustering analysis based on the whole set of CNVs did not show any genetic structure associated with lineages (**Fig.7B**). The high bootstrap support for the first node (100) indicated the existence of two genetic clusters encompassing various sites from the three lineages (**Fig.7B**). Although bootstrap support of the shallower nodes was sometimes low, the classification suggested that geographical proximity and thermal similarity influenced the observed genetic structure (**Fig.7B**). Indeed, NWA sites belonging to genetic cluster 1 (L2, L10, and L6) and 2 (S1, S2, and S3) grouped on the same branch of the dendrogram. Furthermore, geographically close sites of the GRE lineage (G1 and G1, G3 and G4) were also positioned on the same branch. By contrast, the hierarchical analysis based on the 175 candidate CNVs associated with lineage divergence accurately classified the sites of the three lineages (**Fig.7B**). Those results clearly showed that geography and environmental variation (especially temperature in NWA) within the distribution range of each lineage are associated with broad variation in copy number that overrides the genetic signal driven by lineage divergence previously observed with SNP markers (Cayuela et al. 2019). In this previous study, SNPs showed strong genetic differentiation among lineages and very limited genetic structure within lineages.

## Discussion

Our findings support our working hypothesis that CNVs play functional roles pertaining to fitness-related traits, local adaptation and reproductive isolation, in addition to underlying fine-scale genetic structure in capelin from the north Atlantic. We found associations between CNVs and the gonadosomatic index, suggesting that CNVs could affect female fitness by modulating oocyte production and thus fecundity. Second, we detected 105 candidate CNVs associated with temperature, of which 28 (20%) corresponded to genomic regions located within sequences of protein-coding genes. For all candidates, the normalized read depth was negatively correlated with temperature, supporting the hypothesis that fish using “cold water” spawning habitats have more gene copies than their counterparts from warmer spawning habitats. Third, we discovered 175 CNVs associated with the divergence of glacial lineages. We found that 17 to 33% of the candidate CNVs were located within sequences of protein-coding genes, which might lead to the accumulation of genetic incompatibilities and thus could at least partly contribute to reproductive isolation between lineages. Furthermore, we detected an excess of TEs (especially retrotransposons) in candidate CNVs, suggesting that these structural variants have rapidly accumulated during the lineage divergence process. Lastly, clustering analyses revealed genetic structure within capelin lineages resulting from the effects of geography and temperature on local variation in copy numbers. Overall, our results underscore the importance of considering CNVs in population genomics studies, as further discussed below.

### HDplot, a robust approach for CNV discovery in non-model species

In this study, we used HDplot, an approach developed by McKinney et al. (2017) as modified by Dorant et al. (2020). Simulations and empirical data by McKinney et al. (2017) demonstrated the robustness of this approach using a set of simple summary statistics for SNP data by showing that it correctly identified duplicated sequences with >95% concordance for loci of known copy number. Moreover, Dorant et al. (2020) showed that normalized read depth is a reliable proxy for the number of copies of a given duplicated locus. Admittedly, this approach has the same technical limits as RAD-sequencing (i.e., reduced representation of the genome) and other methods of CNV detection based on SNP arrays and whole-genome sequencing (e.g. mappability issues, GC content, PCR duplicates, DNA library quality; Teo et al. 2012). Nevertheless, it provides a useful tool for improving our knowledge on the evolutionary role of duplicated regions and TEs in non-model species by permitting studies involving hundreds of individuals or more from dozens of populations.

### CNVs as a potential determinant of fitness variation

Our study revealed associations between the gonadosomatic index and six candidate CNVs, of which one was positioned within a coding gene with unknown functions. Interestingly, another candidate was found within an intergenic region of ~30 Kbp of chromosome 11 at distance of ~20 Kbp of the gene regulating the receptor of the FSH, a glycoprotein hormone secreted by the pituitary that stimulates early phases of gametogenesis. In female fish, FSH regulates both the secretion of estradiol and the incorporation of vitellogenins into the oocytes (Yaron & Levavi-Sivan 2011), and thus determines the number, the size, and the energy content of eggs that a female can produce. This hormone could therefore be contributing to variation in the gonadosomatic index. The gonadosomatic index was negatively correlated with the normalized read depth of this candidate CNV, suggesting that a high number of copies of the gene (or its promotor) regulating the receptor of the FSH could decrease female fecundity. This duplicated region is thus an interesting candidate potentially involved in the endocrine control of a major component of fitness in capelin and perhaps other fishes.

### CNVs as a potential mechanism promoting thermal adaptation

Our study highlighted CNV-temperature associations, suggesting a possible contribution of these structural variants in thermal adaptation. A total of 105 candidate CNVs were detected using a conservative approach combining regression and redundancy analyses. Twenty-eight of these candidates were located within sequences of protein-coding genes involved in various cellular and molecular processes and could therefore affect the expression of those genes if the duplicated DNA segment includes exons (Kondrashov 2012, Qian & Zhang 2014). Moreover, 72 of the candidate CNVs were located within intergenic regions and could also influence gene expression if they include regulatory regions or belong to large duplicated DNA fragments encompassing one or more genes (Kondrashov 2012, Qian & Zhang 2014). Importantly, the normalized read depth of all the duplicated regions was negatively correlated with temperature, indicating that capelin populations associated with colder waters tend to be characterised by a higher number of copies in various regions of the genome. This result is congruent with a previous study showing that gene duplication is involved in adaptation to cold waters in other (polar) marine fishes (Chen et al. 2008). In this study, Chen et al. (2008) compared the levels of gene expression between the Antarctic toothfish and its warm water dwelling relatives and identified 177 genes with substantial overexpression in the Antarctic toothfish. Many of those genes were related to survival and growth in cold water, and 118 were found to be duplicated, some of them hundreds of times (Chen et al. 2008).

We also showed that five other candidate CNVs were TEs (both DNA transposons and retrotransposons Gypsy LTR) whose normalized read depth, a proxy for their accumulation in the genome, was negatively correlated with temperature. This result is congruent with the hypothesis that TEs can be reactivated in response to external stress, which may favor local adaptation under several environmental conditions (McClintock 1950, Chuong et al. 2017). Thermal stress may indeed cause the derepression of several families of TEs whose activation can in turn induce structural variation, providing a selective advantage under specific climatic conditions (Gonzales et al. 2010). Although this mechanism has been investigated in other organisms such as fruit flies and diatoms (Gonzales et al. 2010, Pargana et al. 2020), it remains largely unexplored in vertebrates. Nevertheless, recent studies suggest that TE accumulation is highly variable among bony fishes (Yuan et al. 2018, Shao et al. 2019). Moreover, a phylogenetic analysis performed on 39 teleost species highlighted an unexpected clusterization of REX3 retroelements isolated from species living in cold waters compared with those isolated from species living in warmer waters, suggesting a possible selective role of temperature on this specific TE (Carducci et al. 2019). In our study system, the accumulation of TEs in the genome of individuals from colder waters could also possibly result from thermal stresses. Alternatively, the variation observed at the population level in copy number of TEs between cold and warm environments could be the outcome of selective processes, such as the one we proposed above for non-TE CNVs associated with temperature. However, the GBS data provided in this study are not sufficient to formulate robust conclusions concerning the functional of role of TEs. This issue could be better addressed through common garden experiments to investigate the influence of thermal stress on TE activation and its consequences on individual fitness (e.g. mortality rate) measured during early developmental stages (e.g., embryo and larvae).

### CNVs as potential drivers of reproductive isolation among nascent species

Our results suggest that CNVs could also be involved in the ongoing speciation process of capelin lineages by enhancing the rate of lineage divergence via the duplication or deletion of genomic regions. Indeed, the accumulation rate of CNVs is 1.5 to 2.5 times higher than that of SNPs in vertebrates (Sudmant et al. 2013, Paudel et al. 2015). Thus, candidate CNVs associated with lineage differentiation may have played a major role in the rapid accumulation of genetic incompatibilities. Although we did not detect any gene enrichment, we showed that 17 to 33% of the candidate CNVs are located inside the sequences of protein-coding genes and could therefore lead to the alteration of gene expression via dosage effects, the disruption of genic functions, and the neofunctionalization of genes (Kondrashov 2012, Qian & Zhang 2014). In turn, those changes could have contributed to the emergence of reproductive barriers, possibly early in the divergence process of glacial lineages.

Our analyses also revealed that among the candidate CNVs, retrotransposons were three to four times more abundant than expected by chance in explaining the differentiation between the three glacial lineages of capelin from the north Atlantic. In contrast, DNA transposons were found in excess for the lineage pair NWA-GRE only. This result strongly suggests that TEs contribute to the structural differentiation of genomes among capelin lineages. Here, it is noteworthy that retrotransposons are expected to accumulate faster in terms of copy number compared to DNA transposons based on their respective mode of transposition (i.e., *copy and paste* for retrotransposons vs. *cut and paste* for DNA transposons) (Calos & Miller 1980). These observations raise the hypothesis that the accumulation of retrotransposons could have contributed to the emergence of reproductive barriers, which could at least partly explain the apparent lack of admixture among capelin lineages in absence of any physical or distance barrier between them (Cayuela et al. 2019).

### CNVs as DNA markers to resolve local adaptive structure

In comparison with our previous study documenting genetic variation in the same species and the same geographic area (Cayuela et al. 2019), our results revealed that CNVs and SNPs show very contrasted patterns of spatial structure, likely due to the different processes underlying their respective evolution. SNP analyses highlighted a pronounced genetic divergence among the three glacial lineages that diverged between 1.8 and 3.5 Mya and a weak intra-lineage genetic structure (Cayuela et al. 2019). This divergence resulted from the high differentiation of ~3000 SNPs (34 to 107 were fixed), likely due to the combined effects of genetic drift, selection and reproductive isolation between lineages (Cayuela et al. 2019). At the intra-lineage level, very large *N_e_* and high gene flow among spawning sites weakens genetic structure, which was only driven by a large-sized chromosomal rearrangement in the NWA lineage (Cayuela et al. 2019).

CNVs showed the opposite pattern: the intra-lineage differentiation exceeded the inter-lineage structure. At the inter-lineage level, divergence is associated with 175 CNVs that likely differentiated during the allopatric phase of geographic isolation. However, the signal of lineage divergence is erased by the effects of geography and environment (especially temperature) which are strongly associated with the variation in copy number within lineages. This pattern could result from the fact that CNVs have a higher mutation rate than SNPs (Redon et al. 2006, Sudmant et al. 2013, Paudel et al. 2015), which may favor rapid evolution under novel environmental conditions (Kondrashov 2012, Qian & Zhang 2014). Rapid variation can also be induced by environmental stresses in the case of TEs (McClintock 1950, Chuong et al. 2017).

Our data suggest that CNV characteristics make them more prone to reveal fine-scale adaptative structure compared to SNPs in a biological system characterised by high gene flow, which is congruent with the conclusions of a previous study in the American lobster (Dorant et al. 2020). Indeed, we showed that capelin experiencing colder waters of Labrador and the Atlantic coast of Newfoundland (cluster 1) were generally characterised by a high number of copies for all CNVs associated with temperature, which could give them a selective advantage in colder environments (Chen et al. 2008). By contrast, individuals inhabiting the warmer waters of the Gulf of St. Lawrence (cluster 2) were systematically characterised by a lower copy number than their counterparts in cluster 1, resulting in pronounced genetic structure within the NWA lineage. Furthermore, the occurrence of complex substructure (i.e., clustering of spatially close spawning sites) within both genetic clusters suggest that other environmental or geographic factors could cause additional local variation in copy number.

The robustness of our analyses based on CNVs is further supported by the unique case of the Saguenay fjord population, which reproduces on beach-spawning sites (S8 in our study) in the St. Lawrence R. estuary, a few kilometers from the mouth of the fjord (Colbeck et al. 2011, Kenchington et al. 2015). The clustering analysis showed that S8 grouped with the cold spawning sites of cluster 1 despite being spatially proximal to sites in cluster 2. This intriguing result is congruent with those of two previous studies (using microsatellite and AFLP markers; Colbeck et al. 2011, Kenchington et al. 2015) showing that the individuals from the Saguenay fjord are more genetically related to those from the Labrador and the northern part of the Newfoundland island (cluster 1) than to those of the Gulf of St. Lawrence (cluster 2). The Saguenay fjord, where the capelin population likely completes its life cycle outside of the breeding period (Lazartigues et al. 2016), is deep (max. ~250 m), has low bottom temperature (between −2 and 1.5 °C. depending on the place and the season, Galbraith et al. 2018), and is a refuge for many Arctic and sub-Arctic species of fish and invertebrates (Drainville 1970, Judkins & Wright 1974). Our results support the hypothesis formulated by Colbeck et al. (2011) and Kenchington et al. (2015) stating that the Saguenay fjord population could be locally adapted to cold waters. Our analyses showed that those individuals have a higher number of copies (for 104 CNVs of the 105 candidates associated with temperature) than those reproducing in the neighboring beach-spawning sites in the Gulf of St. Lawrence. This could due to the retention of cold-adapted individuals in the fjord following the last glacial period.

### Inferring genetic structure using CNVs: research avenues

The study of Dorant et al. (2020) and our work showed that CNVs discovered from RAD-seq data represent an under-explored type of DNA marker that can be investigated in population genomics studies of non-model species and using large datasets. Our analyses unveiled complex hierarchical patterns of structure determined by temperature and local geography that differ from patterns based on SNPs. These two types of markers thus appear highly complementary to document genetic structure, particularly in systems with large *N_e_* and high migration rates, which is the case for the majority of marine species. In these systems, CNVs may be better suited to resolving fine-scale spatial structure driven by contemporary evolutionary processes than SNPs, which could be more efficient to capture large-scale structure resulting from historical and demographic processes. This is consistent with recent studies showing that duplications showed more population-specific structure than SNPs or deletions in humans (Sudmant 2015, Yang 2018) or that revealed pronounced copy-number variation within domestic species (Serres-Armero et al. 2017). With increasing possibilities to investigate all types of genetic variation, not only SNPs but also CNVs or other kinds of SVs, we envision that our approach based on the pioneering work of McKinney al. (2017) will be extended to investigate a broader range of organisms occupying various habitats (i.e., freshwater and terrestrial environments) and characterized by a broad range of demographic characteristics (i.e., small *N_e_*, reduced migration). This would allow further investigation of the environmental and demographic circumstances under which SVs provide a different understanding of population genetic structure compared to SNPs.

## Author contributions

H.C. made the statistical analyses and wrote the paper. Y.D. and E.N. contributed to the bioinformatics and statistical analyses. L.B. and P.S. initiated the project, and L.B. conceptualized and coordinated the work. S.G.-H. performed gonadosomatic index measurements. C.M. and M.L. contributed to the writing of the article. All authors read and edited the final manuscript version.

## Data accessibility

The GBS sequence data used in the analyses will be deposited in DRYAD: ####.

