## Supplementary_material for "Thermal adaptation rather than demographic history drives genetic structure inferred by copy number variants in a marine fish"

**Supplementary material S1**

**Table S1**. Characteristics of the 35 spawning sites considered in the study.

| Site_ID | Lineage | Latitude | Longitude | Type |
| --- | --- | --- | --- | --- |
| P1 | ARC | 66.1148 | 65.73127 | Beach |
| P2 | ARC | 66.14986 | -65.70584 | Beach |
| P3 | ARC | 66.1148 | 65.73127 | Beach |
| G1 | GRE | 69.2494 | -53.5225 | Offshore |
| G2 | GRE | 64.3011 | -51.1833 | Offshore |
| G3 | GRE | 70.458215 | -50.9787 | Offshore |
| G4 | GRE | 64.733333 | -51.1417 | Offshore |
| K1 | NWA | 58.451653 | -62.79610382 | Beach |
| L1 | NWA | 55.12708 | -59.1085 | Demersal |
| L10 | NWA | 51.5258 | -56.8336 | Beach |
| L11 | NWA | 52.3692 | -55.6845 | Demersal |
| L12 | NWA | 51.3830167 | -56.578366 | Beach |
| L13 | NWA | 51.424419 | -57.129263 | Offshore |
| L14 | NWA | 49.482009 | -58.125338 | Beach |
| L15 | NWA | 51.424419 | -57.129263 | Beach |
| L16 | NWA | 51.526 | -56.8346 | Offshore |
| L2 | NWA | 55.0862 | -59.1764 | Beach |
| L3 | NWA | 55.451667 | -57.756667 | Offshore |
| L4 | NWA | 54.9082 | -59.7725 | Demersal |
| L5 | NWA | 54.1799 | -58.4288 | Beach |
| L6 | NWA | 53.4694 | -55.785 | Beach |
| L7 | NWA | 52.7715 | -56.1193 | Demersal |
| L8 | NWA | 52.558677 | -55.766736 | Beach |
| L9 | NWA | 51.6286 | -56.7071 | Demersal |
| N2 | NWA | 47.38165 | -53.46462 | Beach |
| N3 | NWA | 47.40355 | -53.48096 | Demersal |
| N4 | NWA | 47.650694 | -52.696313 | Beach |
| S1 | NWA | 50.266907 | -64.140275 | Beach |
| S2 | NWA | 48.984758 | -64.368145 | Beach |
| S3 | NWA | 48.251333 | -64.75547 | Beach |
| S4 | NWA | 49.21850327 | -68.12046458 | Beach |
| S5 | NWA | 48.78555556 | -67.700833 | Beach |
| S6 | NWA | 48.31732 | -68.871665 | Beach |
| S7 | NWA | 48.22888 | -69.67277777 | Offshore |
| S8 | NWA | 47.576732 | -70.196932 | Beach |

**Table S2.** Repeated DNA elements among the 6620 CNVs detected in our study.

| Transposable_elements | Nb_cnv | Type of repeated elements |
| --- | --- | --- |
| DNA/hAT | 1 | DNA transposon class II |
| DNA/Kolobok-T2 | 1 | DNA transposon class II |
| LINE/L1-Tx1 | 1 | Retrotransposon Non-LTR (class I) |
| LINE/R2-Hero | 1 | Retrotransposon Non-LTR (class I) |
| Satellite | 1 | Satellite DNA |
| DNA/hAT | 2 | DNA transposon class II |
| DNA/hAT-hAT5 | 2 | DNA transposon class II |
| DNA/PIF-Harbinger | 2 | DNA transposon class II |
| DNA/PIF-ISL2EU | 2 | DNA transposon class II |
| DNA/TcMar-ISRm11 | 2 | DNA transposon class II |
| RC/Helitron | 2 | DNA transposon class II |
| LTR/Ngaro | 2 | Retrotransposon LTR (class I) |
| LINE/L1 | 2 | Retrotransposon Non-LTR (class I) |
| DNA/hAT-Charlie | 3 | DNA transposon class II |
| LINE/RTE-BovB | 3 | Retrotransposon Non-LTR (class I) |
| DNA/Crypton-V | 4 | DNA transposon class II |
| DNA/Zisupton | 4 | DNA transposon class II |
| DNA/hAT-Ac | 5 | DNA transposon class II |
| DNA/hAT-Tip100 | 5 | DNA transposon class II |
| DNA/CMC-EnSpm | 6 | DNA transposon class II |
| DNA/Dada | 8 | DNA transposon class II |
| Low_complexity | 11 | Low complexity regions |
| DNA/TcMar-Tc1 | 12 | DNA transposon class II |
| DNA | 14 | DNA transposon class II |
| tRNA | 15 | Transfer RNA |
| LTR/Pao | 16 | Retrotransposon LTR (class I) |
| LINE/L2 | 21 | Retrotransposon Non-LTR (class I) |
| DNA/IS3EU | 22 | DNA transposon class II |
| LTR/Gypsy | 28 | Retrotransposon LTR (class I) |
| LINE/Rex-Babar | 40 | Retrotransposon Non-LTR (class I) |
| Simple_repeat | 89 | Microsatellites |

**Table S3**. LMM outputs for the 105 candidate CNVs detected for temperature. We provide slope coefficient (Slope), the limits of the 95% confidence interval (CI_2.5 and CI_97.5), the marginal R² of the model (corresponding to the amount of variation of normalized read depth explained by temperature), the outputs of a likelihood ratio test ($\chi^{2}$ and Pval), and the FDR value. We also provide the type of genomic region where the CNV is located: coding regions, transposable elements in non-coding regions (DNA transposons and retrotransposons), and other non-coding regions.

| CNV_ID | Slope | CI_2.5 | CI_97.5 | R² | $\chi^{2}$ | Pval | FDR | Type |
| --- | --- | --- | --- | --- | --- | --- | --- | --- |
| 44782_52 | -0.27 | -0.45 | -0.10 | 0.15 | 8.24 | 0.00 | 0.09 | Coding region |
| 46678_68 | -0.26 | -0.47 | -0.05 | 0.12 | 5.70 | 0.02 | 0.09 | Coding region |
| 14190_64 | -0.25 | -0.41 | -0.09 | 0.16 | 8.57 | 0.00 | 0.09 | Coding region |
| 9860_64 | -0.25 | -0.41 | -0.09 | 0.15 | 8.25 | 0.00 | 0.09 | Coding region |
| 62914_54 | -0.25 | -0.41 | -0.08 | 0.14 | 7.79 | 0.01 | 0.09 | Coding region |
| 20807_11 | -0.24 | -0.39 | -0.09 | 0.16 | 8.48 | 0.00 | 0.09 | Coding region |
| 14715_74 | -0.24 | -0.40 | -0.08 | 0.14 | 7.53 | 0.01 | 0.09 | Coding region |
| 17649_32 | -0.24 | -0.39 | -0.08 | 0.15 | 7.92 | 0.00 | 0.09 | Coding region |
| 3183_6 | -0.23 | -0.40 | -0.07 | 0.13 | 7.01 | 0.01 | 0.09 | Coding region |
| 57218_37 | -0.23 | -0.37 | -0.09 | 0.14 | 8.97 | 0.00 | 0.09 | Coding region |
| 24613_12 | -0.23 | -0.38 | -0.07 | 0.12 | 7.45 | 0.01 | 0.09 | Coding region |
| 52786_50 | -0.22 | -0.38 | -0.07 | 0.15 | 7.67 | 0.01 | 0.09 | Coding region |
| 28990_56 | -0.22 | -0.38 | -0.07 | 0.14 | 7.53 | 0.01 | 0.09 | Coding region |
| 58866_42 | -0.22 | -0.37 | -0.07 | 0.14 | 7.64 | 0.01 | 0.09 | Coding region |
| 36851_9 | -0.22 | -0.36 | -0.09 | 0.15 | 9.02 | 0.00 | 0.09 | Coding region |
| 30694_77 | -0.22 | -0.38 | -0.06 | 0.13 | 6.81 | 0.01 | 0.09 | Coding region |
| 19781_57 | -0.22 | -0.36 | -0.08 | 0.14 | 8.29 | 0.00 | 0.09 | Coding region |
| 45721_41 | -0.22 | -0.34 | -0.10 | 0.15 | 10.35 | 0.00 | 0.09 | Coding region |
| 24697_19 | -0.22 | -0.38 | -0.05 | 0.11 | 6.39 | 0.01 | 0.09 | Coding region |
| 13380_75 | -0.21 | -0.37 | -0.05 | 0.12 | 6.19 | 0.01 | 0.09 | Coding region |
| 17635_74 | -0.21 | -0.33 | -0.08 | 0.13 | 8.61 | 0.00 | 0.09 | Coding region |
| 3_62 | -0.20 | -0.33 | -0.07 | 0.14 | 8.26 | 0.00 | 0.09 | Coding region |
| 5326_9 | -0.20 | -0.36 | -0.05 | 0.12 | 6.30 | 0.01 | 0.09 | Coding region |
| 62973_22 | -0.20 | -0.35 | -0.05 | 0.12 | 6.50 | 0.01 | 0.09 | Coding region |
| 377_68 | -0.20 | -0.33 | -0.07 | 0.12 | 7.84 | 0.01 | 0.09 | Coding region |
| 45979_12 | -0.18 | -0.30 | -0.06 | 0.13 | 7.67 | 0.01 | 0.09 | Coding region |
| 4231_24 | -0.18 | -0.29 | -0.07 | 0.13 | 8.57 | 0.00 | 0.09 | Coding region |
| 18625_22 | -0.17 | -0.27 | -0.07 | 0.13 | 9.51 | 0.00 | 0.09 | Coding region |
| 4669_61 | -0.35 | -0.57 | -0.13 | 0.19 | 8.54 | 0.00 | 0.09 | DNA transposon |
| 11762_29 | -0.31 | -0.54 | -0.09 | 0.15 | 7.05 | 0.01 | 0.09 | Retrotransposon |
| 45132_27 | -0.25 | -0.43 | -0.07 | 0.12 | 6.85 | 0.01 | 0.09 | DNA transposon |
| 18013_49 | -0.21 | -0.35 | -0.07 | 0.14 | 8.15 | 0.00 | 0.09 | DNA transposon |
| 27856_29 | -0.20 | -0.35 | -0.05 | 0.12 | 6.52 | 0.01 | 0.09 | DNA transposon |
| 54506_10 | -0.78 | -1.32 | -0.24 | 0.22 | 7.28 | 0.01 | 0.09 | Non-coding region |
| 43429_52 | -0.62 | -1.05 | -0.18 | 0.17 | 7.00 | 0.01 | 0.09 | Non-coding region |
| 29554_79 | -0.34 | -0.61 | -0.06 | 0.12 | 5.61 | 0.02 | 0.09 | Non-coding region |
| 35447_51 | -0.36 | -0.61 | -0.11 | 0.15 | 7.12 | 0.01 | 0.09 | Non-coding region |
| 40484_54 | -0.31 | -0.53 | -0.09 | 0.15 | 7.11 | 0.01 | 0.09 | Non-coding region |
| 42417_41 | -0.29 | -0.48 | -0.10 | 0.13 | 8.15 | 0.00 | 0.09 | Non-coding region |
| 57247_54 | -0.27 | -0.45 | -0.09 | 0.15 | 7.56 | 0.01 | 0.09 | Non-coding region |
| 43266_6 | -0.27 | -0.45 | -0.10 | 0.18 | 8.43 | 0.00 | 0.09 | Non-coding region |
| 25048_60 | -0.26 | -0.44 | -0.09 | 0.18 | 7.81 | 0.01 | 0.09 | Non-coding region |
| 14986_19 | -0.27 | -0.43 | -0.10 | 0.17 | 8.74 | 0.00 | 0.09 | Non-coding region |
| 50035_59 | -0.25 | -0.43 | -0.08 | 0.14 | 7.47 | 0.01 | 0.09 | Non-coding region |
| 20996_76 | -0.24 | -0.42 | -0.07 | 0.13 | 6.77 | 0.01 | 0.09 | Non-coding region |
| 31220_53 | -0.24 | -0.42 | -0.07 | 0.14 | 6.90 | 0.01 | 0.09 | Non-coding region |
| 51780_62 | -0.25 | -0.41 | -0.09 | 0.14 | 7.91 | 0.00 | 0.09 | Non-coding region |
| 57524_53 | -0.23 | -0.41 | -0.05 | 0.11 | 5.93 | 0.01 | 0.09 | Non-coding region |
| 34287_79 | -0.23 | -0.41 | -0.05 | 0.12 | 5.85 | 0.02 | 0.09 | Non-coding region |
| 20012_43 | -0.23 | -0.40 | -0.06 | 0.12 | 6.64 | 0.01 | 0.09 | Non-coding region |
| 12742_58 | -0.25 | -0.40 | -0.10 | 0.19 | 9.49 | 0.00 | 0.09 | Non-coding region |
| 43507_50 | -0.23 | -0.39 | -0.07 | 0.12 | 7.01 | 0.01 | 0.09 | Non-coding region |
| 49388_7 | -0.24 | -0.39 | -0.08 | 0.13 | 7.99 | 0.00 | 0.09 | Non-coding region |
| 35049_76 | -0.23 | -0.39 | -0.07 | 0.13 | 7.09 | 0.01 | 0.09 | Non-coding region |
| 51928_39 | -0.23 | -0.39 | -0.08 | 0.15 | 7.70 | 0.01 | 0.09 | Non-coding region |
| 45973_78 | -0.23 | -0.38 | -0.09 | 0.15 | 8.44 | 0.00 | 0.09 | Non-coding region |
| 36654_66 | -0.22 | -0.38 | -0.07 | 0.12 | 7.19 | 0.01 | 0.09 | Non-coding region |
| 58255_54 | -0.22 | -0.38 | -0.06 | 0.13 | 7.03 | 0.01 | 0.09 | Non-coding region |
| 38615_31 | -0.24 | -0.38 | -0.10 | 0.16 | 9.20 | 0.00 | 0.09 | Non-coding region |
| 42785_44 | -0.22 | -0.38 | -0.07 | 0.12 | 7.09 | 0.01 | 0.09 | Non-coding region |
| 22083_31 | -0.23 | -0.38 | -0.08 | 0.14 | 8.34 | 0.00 | 0.09 | Non-coding region |
| 13116_63 | -0.21 | -0.38 | -0.05 | 0.12 | 5.95 | 0.01 | 0.09 | Non-coding region |
| 26395_6 | -0.22 | -0.38 | -0.07 | 0.13 | 7.43 | 0.01 | 0.09 | Non-coding region |
| 54868_77 | -0.24 | -0.38 | -0.11 | 0.17 | 10.72 | 0.00 | 0.09 | Non-coding region |
| 26097_75 | -0.23 | -0.38 | -0.09 | 0.16 | 8.82 | 0.00 | 0.09 | Non-coding region |
| 8041_56 | -0.23 | -0.38 | -0.09 | 0.16 | 9.03 | 0.00 | 0.09 | Non-coding region |
| 33977_9 | -0.23 | -0.37 | -0.08 | 0.14 | 7.91 | 0.00 | 0.09 | Non-coding region |
| 56181_28 | -0.21 | -0.37 | -0.05 | 0.11 | 6.29 | 0.01 | 0.09 | Non-coding region |
| 40393_57 | -0.23 | -0.37 | -0.08 | 0.14 | 8.21 | 0.00 | 0.09 | Non-coding region |
| 36783_53 | -0.22 | -0.37 | -0.06 | 0.14 | 6.96 | 0.01 | 0.09 | Non-coding region |
| 9148_5 | -0.22 | -0.37 | -0.07 | 0.13 | 7.29 | 0.01 | 0.09 | Non-coding region |
| 27711_54 | -0.22 | -0.37 | -0.07 | 0.15 | 7.85 | 0.01 | 0.09 | Non-coding region |
| 30954_6 | -0.23 | -0.37 | -0.10 | 0.15 | 9.48 | 0.00 | 0.09 | Non-coding region |
| 38700_21 | -0.22 | -0.37 | -0.08 | 0.13 | 7.93 | 0.00 | 0.09 | Non-coding region |
| 55807_60 | -0.21 | -0.36 | -0.06 | 0.12 | 7.03 | 0.01 | 0.09 | Non-coding region |
| 2333_19 | -0.22 | -0.36 | -0.08 | 0.13 | 8.28 | 0.00 | 0.09 | Non-coding region |
| 23974_73 | -0.21 | -0.36 | -0.06 | 0.12 | 6.92 | 0.01 | 0.09 | Non-coding region |
| 50661_68 | -0.22 | -0.36 | -0.07 | 0.13 | 7.91 | 0.00 | 0.09 | Non-coding region |
| 38792_8 | -0.22 | -0.36 | -0.09 | 0.14 | 8.96 | 0.00 | 0.09 | Non-coding region |
| 36903_16 | -0.21 | -0.35 | -0.08 | 0.15 | 8.95 | 0.00 | 0.09 | Non-coding region |
| 5183_12 | -0.22 | -0.35 | -0.10 | 0.17 | 10.10 | 0.00 | 0.09 | Non-coding region |
| 10876_27 | -0.20 | -0.34 | -0.06 | 0.14 | 7.33 | 0.01 | 0.09 | Non-coding region |
| 41625_15 | -0.20 | -0.34 | -0.05 | 0.13 | 6.75 | 0.01 | 0.09 | Non-coding region |
| 3688_75 | -0.20 | -0.34 | -0.07 | 0.13 | 7.67 | 0.01 | 0.09 | Non-coding region |
| 29035_60 | -0.20 | -0.34 | -0.05 | 0.12 | 6.81 | 0.01 | 0.09 | Non-coding region |
| 44926_10 | -0.23 | -0.34 | -0.12 | 0.14 | 13.35 | 0.00 | 0.09 | Non-coding region |
| 42338_39 | -0.19 | -0.34 | -0.04 | 0.11 | 5.90 | 0.02 | 0.09 | Non-coding region |
| 44879_18 | -0.21 | -0.33 | -0.08 | 0.15 | 9.04 | 0.00 | 0.09 | Non-coding region |
| 28679_68 | -0.19 | -0.33 | -0.06 | 0.13 | 7.09 | 0.01 | 0.09 | Non-coding region |
| 12597_77 | -0.19 | -0.33 | -0.06 | 0.13 | 6.97 | 0.01 | 0.09 | Non-coding region |
| 640_11 | -0.21 | -0.33 | -0.09 | 0.16 | 9.68 | 0.00 | 0.09 | Non-coding region |
| 13190_5 | -0.19 | -0.33 | -0.05 | 0.12 | 6.80 | 0.01 | 0.09 | Non-coding region |
| 56523_71 | -0.20 | -0.33 | -0.07 | 0.14 | 8.04 | 0.00 | 0.09 | Non-coding region |
| 2958_41 | -0.20 | -0.32 | -0.08 | 0.14 | 8.94 | 0.00 | 0.09 | Non-coding region |
| 55476_6 | -0.19 | -0.32 | -0.06 | 0.12 | 7.19 | 0.01 | 0.09 | Non-coding region |
| 47581_44 | -0.19 | -0.32 | -0.06 | 0.14 | 7.28 | 0.01 | 0.09 | Non-coding region |
| 27535_29 | -0.19 | -0.32 | -0.06 | 0.11 | 7.50 | 0.01 | 0.09 | Non-coding region |
| 39977_62 | -0.21 | -0.32 | -0.10 | 0.16 | 11.40 | 0.00 | 0.09 | Non-coding region |
| 13863_75 | -0.20 | -0.31 | -0.08 | 0.14 | 9.10 | 0.00 | 0.09 | Non-coding region |
| 5512_54 | -0.19 | -0.31 | -0.07 | 0.12 | 8.51 | 0.00 | 0.09 | Non-coding region |
| 5787_77 | -0.20 | -0.31 | -0.09 | 0.12 | 10.84 | 0.00 | 0.09 | Non-coding region |
| 6579_65 | -0.20 | -0.31 | -0.08 | 0.15 | 9.30 | 0.00 | 0.09 | Non-coding region |
| 35501_59 | -0.18 | -0.31 | -0.06 | 0.12 | 7.30 | 0.01 | 0.09 | Non-coding region |
| 10343_26 | -0.19 | -0.31 | -0.07 | 0.13 | 8.63 | 0.00 | 0.09 | Non-coding region |
| 32788_74 | -0.18 | -0.30 | -0.06 | 0.13 | 7.68 | 0.01 | 0.09 | Non-coding region |

**Table S4.** CNVs located within the sequence of coding genes associated with lineage divergence. We provide the CNV ID (CNV_ID), the pair of lineages for which the CNV has been detected as a candidate, the gene name in the annotated Capelin reference genome, and the Pfam of the corresponding gene.

| CNV_ID | Gene_ID | Pfam |
| --- | --- | --- |
| 5326_9 | XM_012826588.1 | PF00083 |
| 5326_9 | XM_012827983.1 | PF00028; PF01049; PF08758 |
| 13380_75 | XM_012826816.1 | PF00583 |
| 13380_75 | XM_012838233.1 | PF00069 |
| 18625_22 | XM_012830463.1 | _ |
| 18625_22 | XM_012839307.1 | _ |
| 19781_57 | XM_012831799.1 | _ |
| 19781_57 | XM_012831799.1 | _ |
| 24613_12 | XM_012835216.1 | PF11816;PF00400 |
| 24697_19 | XM_012820953.1 | PF01534;PF01392 |
| 24697_19 | XM_012839981.1 | PF01534;PF01392 |
| 28990_56 | XM_012834723.1 | _ |
| 28990_56 | XM_012818751.1 | PF00224;PF02887 |
| 36851_9 | XM_012827684.1 | PF00335 |
| 44782_52 | XM_012825231.1 | PF00069 |
| 44782_52 | XM_012825205.1 | PF02862 |
| 44782_52 | XM_012828414.1 | PF00041;PF07679 |
| 44782_52 | XM_012821891.1 | PF00651;PF00096 |
| 44782_52 | XM_012839786.1 | PF00011 |
| 44782_52 | XM_012831586.1 | PF00012 |
| 45721_41 | XM_012832030.1 | PF09368;PF04000 |
| 46678_68 | XM_012815663.1 | _ |
| 57218_37 | XM_012840232.1 | PF08264;PF00043;PF00133;PF10458 |
| 58866_42 | XM_012825342.1 | PF00628;PF02182;PF12148;PF00240;PF00097 |
| 58866_42 | XM_012823211.1 | PF02759;PF13901 |
| 58866_42 | XM_012823211.1 | PF02373;PF02375;PF18104 |
| 3_62 | XM_012836231.1 | PF00076 |
| 377_68 | XM_012831660.1 | PF00041;PF07679;PF00102 |
| 377_68 | XM_012831763.1 | PF03114;PF00018;PF00104;PF00105 |
| 3183_6 | XM_012826038.1 | PF10174 |
| 4231_24 | XM_012841316.1 | PF13639 |
| 4231_24 | XM_012825652.1 | _ |
| 4231_24 | XM_012825654.1 | _ |
| 4231_24 | XM_012819084.1 | PF06367;PF06371;PF02181 |
| 4231_24 | XM_012825650.1 | _ |
| 4231_24 | XM_012825683.1 | PF00025 |
| 9860_64 | XM_012834758.1 | PF02174;PF00169 |
| 14190_64 | XM_012834574.1 | PF00027;PF02197 |
| 14715_74 | XM_012821880.1 | PF13639 |
| 14715_74 | XM_012826610.1 | PF13639 |
| 17635_74 | XM_012826879.1 | PF00626;PF08033;PF04815;PF04811;PF04810 |
| 17649_32 | XM_012826828.1 | PF01388;PF08169;PF11717 |
| 20807_11 | XM_012829317.1 | PF00168;PF07002 |
| 30694_77 | XM_012820730.1 | _ |
| 30694_77 | XM_012824897.1 | PF00285 |
| 30694_77 | XM_012824873.1 | _ |
| 30694_77 | XM_012824872.1 | _ |
| 45979_12 | XM_012818767.1 | PF00071 |
| 52786_50 | XM_012828307.1 | PF12796;PF03020 |
| 62914_54 | XM_012826553.1 | PF08920 |
| 62914_54 | XM_012826552.1 | PF08920 |
| 62973_22 | XM_012826562.1 | PF07986 |

**Table S5.** CNVs associated with lineage divergence located within the sequence of coding genes. We provide the CNV ID (CNV_ID), the pair of lineages for which the CNV has been detected as a candidate, the gene name in the annotated Capelin reference genome, and the Pfam of the corresponding gene.

| CNV_ID | ARC-NWA | ARC-GRE | NWA-GRE | Gene_ID | Pfam |
| --- | --- | --- | --- | --- | --- |
| 8007_49 | YES | _ | YES | XM_012822075.1 | _ |
| 10019_52 | _ | YES | _ | XM_012841636.1 | PF07443;PF00271;PF00176 |
| 17091_11 | YES | _ | YES | XM_012833286.1 | PF00067 |
| 17091_11 | YES | _ | YES | XM_012833280.1 | PF00067 |
| 17265_72 | YES | _ | YES | XM_012818645.1 | PF02338;PF01754 |
| 17426_73 | _ | _ | YES | XM_012835172.1 | PF02207 |
| 19919_55 | YES | _ | _ | XM_012831799.1 | _ |
| 19919_55 | YES | _ | _ | XM_012831798.1 | _ |
| 19919_55 | YES | _ | _ | XM_012831753.1 | PF00651 |
| 19919_55 | YES | _ | _ | XM_012840298.1 | PF00168;PF00632;PF00397 |
| 19919_55 | YES | _ | _ | XM_012840297.1 | PF00168;PF00632;PF00397 |
| 19919_55 | YES | _ | _ | XM_012840295.1 | PF00168;PF00632;PF00397 |
| 25625_71 | _ | YES | YES | XM_012841639.1 | PF00063;PF00169;PF00621 |
| 25625_71 | _ | YES | YES | XM_012829438.1 | PF00019;PF00688 |
| 30021_73 | YES | YES | _ | XM_012833860.1 | PF11838;PF01433 |
| 32926_48 | _ | YES | _ | XM_012840138.1 | PF01603 |
| 32926_48 | _ | YES | _ | XM_012816361.1 | PF00027;PF00520;PF13426 |
| 34515_40 | YES | _ | YES | XM_012834483.1 | _ |
| 34480_73 | _ | _ | YES | XM_012831519.1 | PF07707;PF00651;PF01344 |
| 40530_24 | _ | _ | YES | XM_012821160.1 | PF15435 |
| 40530_24 | _ | _ | YES | XM_012821189.1 | PF01490 |
| 42014_66 | _ | YES | _ | XM_012823961.1 | _ |
| 43952_28 | _ | YES | _ | XM_012816389.1 | PF00307;PF00681;PF03501 |
| 44685_20 | _ | YES | YES | XM_012837952.1 | PF06911 |
| 44749_14 | YES | YES | YES | XM_012825231.1 | PF00069 |
| 44749_14 | YES | YES | YES | XM_012825205.1 | PF02862 |
| 44749_14 | YES | YES | YES | XM_012831544.1 | _ |
| 44749_14 | YES | YES | YES | XM_012828414.1 | PF00041;PF07679 |
| 44749_14 | YES | YES | _ | XM_012821891.1 | PF00640;PF00397 |
| 44749_14 | YES | YES | _ | XM_012827695.1 | PF07651 |
| 46081_18 | YES | _ | _ | XM_012816662.1 | PF00063;PF02736;PF01576 |
| 46287_30 | YES | _ | YES | XM_012819900.1 | PF12906 |
| 47967_76 | YES | YES | _ | XR_001162481.1 | _ |
| 48678_50 | _ | _ | YES | XM_012816249.1 | PF02985;PF15017 |
| 49215_47 | _ | YES | YES | XM_012822498.1 | PF00125;PF16211 |
| 49215_47 | _ | YES | YES | XM_012839022.1 | PF00125;PF16211 |
| 49215_47 | _ | YES | YES | XM_012829047.1 | PF00125;PF16211 |
| 49215_47 | _ | YES | YES | XM_012823630.1 | PF00125;PF16211 |
| 52083_58 | YES | _ | YES | XM_012837507.1 | PF05694 |
| 52083_58 | YES | _ | YES | XM_012837483.1 | PF00454 |
| 52932_71 | _ | _ | YES | XM_012830002.1 | _ |
| 54934_49 | _ | _ | YES | XM_012827265.1 | PF07714 |
| 54076_5 | YES | YES | _ | XM_012824630.1 | PF01063 |
| 54225_76 | _ | YES | _ | XM_012837112.1 | PF00017;PF07525 |
| 54225_76 | _ | YES | _ | XM_012837140.1 | PF00169;PF01369 |
| 54225_76 | _ | YES | _ | XM_012819066.1 | _ |
| 55180_25 | YES | YES | _ | XM_012819066.1 | PF00675;PF05193;PF16187 |
| 56444_58 | YES | _ | YES | XM_012842344.1 | PF08767 |
| 58377_48 | YES | _ | _ | XM_012835987.1 | PF00646;PF16866;PF02008 |
| 58917_41 | YES | YES | _ | XM_012823211.1 | PF02759;PF13901 |
| 59169_46 | _ | _ | YES | XM_012821436.1 | _ |
| 61048_12 | YES | _ | YES | XM_012839708.1 | PF00002;PF02793 |
| 61048_12 | YES | _ | YES | XM_012839750.1 | PF00595 |
| 61088_71 | YES | _ | YES | XM_012819096.1 | PF04880 |
| 61088_71 | YES | _ | YES | XM_012839735.1 | PF04880 |
| 61088_71 | YES | _ | YES | XM_012819095.1 | PF04880 |
| 2263_72 | _ | YES | _ | XM_012823763.1 | PF00621 |
| 2263_72 | _ | YES | _ | XM_012823739.1 | _ |
| 2671_69 | YES | YES | _ | XM_012815886.1 | _ |
| 3415_52 | YES | YES | _ | XM_012830361.1 | _ |
| 81087_63 | YES | YES | _ | XM_012838229.1 | PF07679;PF07714 |
| 5124_46 | _ | YES | _ | XM_012814283.1 | PF01967;PF06463;PF04055 |
| 6812_58 | YES | _ | _ | XM_012840423.1 | PF00653;PF12356;PF00179 |
| 7611_23 | _ | YES | _ | XM_012823231.1 | PF00096 |
| 9645_33 | YES | YES | _ | XM_012818559.1 | PF08355;PF08356;PF00071 |
| 10641_37 | YES | _ | _ | XM_012815482.1 | PF12796;PF15898 |
| 10641_37 | YES | _ | _ | XM_012815478.1 | PF12796;PF15898 |
| 10685_25 | _ | YES | _ | XM_012823531.1 | PF01167;PF16322 |
| 11217_61 | _ | YES | _ | XM_012834367.1 | PF12894;PF08662;PF00400 |
| 11217_61 | _ | YES | _ | XM_012830412.1 | PF01237;PF00169 |
| 11319_9 | YES | _ | YES | XM_012839247.1 | _ |
| 11951_40 | _ | YES | _ | XM_012815820.1 | PF00038;PF04732 |
| 14514_57 | YES | _ | YES | XM_012840990.1 | PF00069 |
| 15662_71 | _ | YES | _ | XM_012836689.1 | PF02210;PF01034 |
| 15662_71 | _ | YES | _ | XM_012815555.1 | PF02210;PF01034 |
| 16493_76 | _ | YES | _ | XM_012831001.1 | _ |
| 17376_75 | _ | _ | YES | XM_012842408.1 | PF00328;PF18086 |
| 17632_54 | YES | _ | YES | XM_012826879.1 | PF00626;PF08033;PF04815;PF04811;PF04810 |
| 19304_44 | YES | YES | _ | XM_012829804.1 | PF02138;PF14844;PF00400 |
| 19491_61 | YES | _ | _ | XM_012841819.1 | PF00307;PF12510 |
| 20723_78 | _ | YES | YES | XM_012841298.1 | PF13516;PF05729 |
| 21629_64 | _ | _ | YES | XM_012825041.1 | PF04103 |
| 24134_17 | _ | YES | _ | XM_012817390.1 | PF08961;PF17169 |
| 25138_26 | YES | YES | _ | XM_012829097.1 | PF07686 |
| 27942_6 | YES | _ | _ | XM_012820838.1 | PF00046;PF00292;PF12360 |
| 28085_8 | YES | _ | _ | XM_012818274.1 | PF00400 |
| 28652_60 | YES | _ | _ | XM_012832188.1 | PF00995 |
| 29409_70 | _ | YES | _ | XM_012814509.1 | PF03114;PF00621;PF00018;PF07653;PF14604 |
| 29626_70 | YES | YES | _ | XM_012838093.1 | PF10534;PF06663;PF00595;PF00536 |
| 30716_11 | _ | YES | _ | XM_012820730.1 | PF15337 |
| 30716_11 | _ | YES | _ | XR_001162275.1 | PF15337 |
| 30716_11 | _ | YES | _ | XM_012834975.1 | PF15337 |
| 30716_11 | _ | YES | _ | XM_012824875.1 | PF15337 |
| 32815_73 | _ | YES | _ | XM_012841402.1 | PF07724;PF12796;PF10431 |
| 35352_35 | _ | _ | YES | XM_012822579.1 | PF13882;PF00041;PF07679;PF00047 |
| 35707_17 | _ | YES | _ | XM_012814493.1 | PF00002 |
| 35778_21 | _ | YES | _ | XM_012819024.1 | PF00069 |
| 36188_64 | _ | _ | YES | XM_012824547.1 | PF14911;PF14910 |
| 37885_14 | _ | YES | _ | XM_012828958.1 | PF02809;PF13519 |
| 40130_45 | YES | YES | _ | XM_012825386.1 | PF00130;PF00069;PF00433 |
| 40130_45 | YES | YES | _ | XM_012825393.1 | PF01585;PF00076 |
| 41227_61 | _ | YES | _ | XM_012822344.1 | PF00010;PF00989 |
| 42750_74 | YES | _ | _ | XM_012837100.1 | PF08366;PF00400 |
| 43069_35 | _ | YES | _ | XM_012821739.1 | _ |
| 43272_49 | YES | _ | YES | XM_012837119.1 | PF00622 |
| 43272_49 | YES | _ | YES | XM_012837111.1 | PF07690 |
| 44094_54 | _ | _ | YES | XM_012816685.1 | _ |
| 44094_54 | _ | _ | YES | XM_012816666.1 | _ |
| 45578_38 | YES | _ | _ | XM_012830006.1 | PF01576 |
| 46879_61 | YES | YES | YES | XM_012816434.1 | PF15924;PF00534 |
| 50491_69 | YES | YES | _ | XM_012824394.1 | PF00040;PF00059;PF00652 |
| 50962_65 | YES | YES | _ | XM_012837582.1 | PF01419 |
| 52746_50 | YES | _ | _ | XM_012818418.1 | PF00520 |
| 54072_76 | _ | YES | _ | XM_012832532.1 | PF13516;PF05729 |
| 55021_71 | YES | _ | _ | XM_012834865.1 | PF00125;PF16211 |
| 63373_42 | _ | YES | _ | XM_012824399.1 | PF14658;PF14662;PF05781 |
| 62613_11 | _ | YES | _ | XM_012832109.1 | PF00685 |

**Table S6**. Candidate CNVs associated with lineage divergence and corresponding to repeated elements.

| CNV_id | Contig | Type | repeat_Class_familiy |
| --- | --- | --- | --- |
| **ARC-GRE** |  |  |  |
| 15408_46 | contig_21 | Retrotransposon | LINE/Rex-Babar |
| 48334_77 | contig_55 | Retrotransposon | LINE/Rex-Babar |
| 11016_18 | contig_171 | DNA transposon | DNA/CMC-EnSpm |
| 21001_70 | contig_271 | Interspersed repeat | Low_complexity |
| 30021_73 | contig_345 | Retrotransposon | LINE/Rex-Babar |
| 43161_15 | contig_445 | Retrotransposon | LTR/Ngaro |
| 45161_51 | contig_480 | DNA transposon | DNA/Zisupton |
| 50809_53 | contig_620 | Interspersed repeat | Simple_repeat |
| 10924_65 | contig_1696 | Retrotransposon | LINE/Rex-Babar |
| 11113_53 | contig_1734 | Retrotransposon | LTR/Gypsy |
| 14481_22 | contig_2080 | Retrotransposon | LTR/Gypsy |
| 15103_6 | contig_2188 | Retrotransposon | LINE/Rex-Babar |
| 21588_8 | contig_2762 | Retrotransposon | LTR/Gypsy |
| 45580_70 | contig_4944 | DNA transposon | DNA/IS3EU |
| 46772_40 | contig_5238 | DNA transposon | DNA/hAT-Tip100 |
| **ARC-NWA** |  |  |  |
| 45132_27 | contig_47 | DNA transposon | DNA/Dada |
| 51276_52 | contig_63 | Retrotransposon | LTR/Gypsy |
| 10419_16 | contig_162 | Retrotransposon | LTR/Pao |
| 11016_18 | contig_171 | DNA transposon | DNA/CMC-EnSpm |
| 17265_72 | contig_236 | Retrotransposon | LTR/Gypsy |
| 21001_70 | contig_271 | Interspersed repeat | Low_complexity |
| 30021_73 | contig_345 | Retrotransposon | LINE/Rex-Babar |
| 31418_40 | contig_355 | Retrotransposon | LINE/Rex-Babar |
| 11113_53 | contig_1734 | Retrotransposon | LTR/Gypsy |
| 11319_9 | contig_1766 | DNA transposon | DNA/IS3EU |
| 11329_26 | contig_1766 | DNA transposon | DNA/IS3EU |
| 14514_57 | contig_2089 | Interspersed repeat | Simple_repeat |
| 21588_8 | contig_2762 | Retrotransposon | LTR/Gypsy |
| 46776_55 | contig_5238 | Retrotransposon | LTR/Gypsy |
| **NWA-GRE** |  |  |  |
| 15408_46 | contig_21 | Retrotransposon | LINE/Rex-Babar |
| 45132_27 | contig_47 | DNA transposon | DNA/Dada |
| 48334_77 | contig_55 | Retrotransposon | LINE/Rex-Babar |
| 51276_52 | contig_63 | Retrotransposon | LTR/Gypsy |
| 10419_16 | contig_162 | Retrotransposon | LTR/Pao |
| 17265_72 | contig_236 | Retrotransposon | LTR/Gypsy |
| 21001_70 | contig_271 | Interspersed repeat | Low_complexity |
| 23769_15 | contig_296 | DNA transposon | DNA |
| 31418_40 | contig_355 | Retrotransposon | LINE/Rex-Babar |
| 43161_15 | contig_445 | Retrotransposon | LTR/Ngaro |
| 45161_51 | contig_480 | DNA transposon | DNA/Zisupton |
| 59169_46 | contig_887 | Retrotransposon | LTR/Pao |
| 11319_9 | contig_1766 | DNA transposon | DNA/IS3EU |
| 11329_26 | contig_1766 | DNA transposon | DNA/IS3EU |
| 14481_22 | contig_2080 | Retrotransposon | LTR/Gypsy |
| 14514_57 | contig_2089 | Interspersed repeat | Simple_repeat |
| 38406_15 | contig_4022 | DNA transposon | DNA |
| 42930_74 | contig_4355 | Retrotransposon | LINE/Rex-Babar |
| 46772_40 | contig_5238 | DNA transposon | DNA/hAT-Tip100 |
| 49370_38 | contig_5825 | DNA transposon | DNA/IS3EU |
| 54760_19 | contig_7477 | DNA transposon | DNA/Dada |

**Table S7**. Regression outputs of the models where we examined how read depth of the 105 candidate CNVs associated with temperature differed between Saguenay population (S8) and neighboring spawning sites in the St Lawrence estuary (S1, S2, S3, S4, S5, and S6). The identity of spawning sites was coded as a discrete variable with two modalities (S8 vs other sites). We provide the slope coefficient (Slope) for the modality “S8” and the FDR value.

| CNV_ID | Slope | FDR |
| --- | --- | --- |
| 45132_27 | 0.6072538 | 3.09E-09 |
| 58255_54 | 0.79630705 | 2.09E-16 |
| 5326_9 | 0.62870374 | 1.25E-12 |
| 11762_29 | 1.14578893 | 4.25E-18 |
| 12597_77 | 0.5893926 | 9.47E-14 |
| 13380_75 | 0.60102184 | 2.99E-12 |
| 14986_19 | 0.72976137 | 4.83E-16 |
| 18625_22 | 0.26005486 | 0.00039433 |
| 19781_57 | 0.65003866 | 1.54E-12 |
| 20996_76 | 0.87496182 | 3.87E-17 |
| 24613_12 | 0.91187729 | 4.03E-19 |
| 24697_19 | 0.68541809 | 3.73E-12 |
| 25048_60 | 0.93506098 | 3.15E-30 |
| 28990_56 | 0.75413244 | 4.22E-17 |
| 29554_79 | 1.12382696 | 8.20E-16 |
| 36851_9 | 0.63432662 | 1.26E-12 |
| 36903_16 | 0.56194316 | 3.01E-12 |
| 40484_54 | 0.89575322 | 3.06E-13 |
| 42417_41 | 1.01576052 | 8.60E-19 |
| 43429_52 | 2.62025443 | 4.45E-36 |
| 43507_50 | 0.8446455 | 7.36E-17 |
| 44782_52 | 0.84802174 | 4.10E-14 |
| 44879_18 | 0.54133075 | 8.35E-11 |
| 45721_41 | 0.50045471 | 5.18E-09 |
| 46678_68 | 0.92453878 | 1.36E-16 |
| 47581_44 | 0.57412901 | 6.10E-14 |
| 49388_7 | 0.63942369 | 1.09E-09 |
| 50661_68 | 0.61934025 | 1.17E-10 |
| 51780_62 | 0.81687588 | 3.18E-15 |
| 51928_39 | 0.55401939 | 3.76E-11 |
| 54868_77 | 0.53227525 | 3.24E-11 |
| 56181_28 | 0.72714616 | 2.03E-12 |
| 56523_71 | 0.43961681 | 3.02E-09 |
| 57218_37 | 0.63347483 | 1.49E-11 |
| 57247_54 | 0.83803145 | 1.39E-15 |
| 57524_53 | 0.72704426 | 3.05E-12 |
| 58866_42 | 0.66390226 | 3.10E-14 |
| 3_62 | 0.6399182 | 9.20E-14 |
| 377_68 | 0.55694352 | 1.68E-10 |
| 640_11 | 0.55101935 | 7.61E-13 |
| 2333_19 | 0.69154469 | 4.47E-15 |
| 2958_41 | 0.28192337 | 0.00019151 |
| 3183_6 | 0.77381206 | 8.40E-14 |
| 3688_75 | 0.47979368 | 2.09E-09 |
| 4231_24 | 0.43060735 | 5.68E-09 |
| 4669_61 | 1.2112878 | 4.18E-31 |
| 5183_12 | 0.63851349 | 6.22E-17 |
| 5512_54 | 0.41054051 | 1.58E-06 |
| 5787_77 | 0.36374573 | 7.88E-05 |
| 6579_65 | 0.52260459 | 1.72E-12 |
| 8041_56 | 0.76900403 | 8.13E-19 |
| 9148_5 | 0.67387391 | 9.76E-14 |
| 9860_64 | 0.79582382 | 6.35E-16 |
| 10343_26 | 0.52260805 | 1.87E-11 |
| 10876_27 | 0.56574341 | 2.59E-13 |
| 12742_58 | 0.62811261 | 4.98E-14 |
| 13116_63 | 0.72020253 | 2.47E-15 |
| 13190_5 | 0.58006097 | 2.05E-11 |
| 13863_75 | 0.57278162 | 9.11E-12 |
| 14190_64 | 0.82947072 | 8.90E-19 |
| 14715_74 | 0.76615602 | 5.14E-15 |
| 17635_74 | 0.53365465 | 1.15E-09 |
| 17649_32 | 0.68486251 | 4.96E-13 |
| 18013_49 | 0.65004166 | 6.85E-13 |
| 20012_43 | 0.82962571 | 2.79E-16 |
| 20807_11 | 0.62254832 | 9.71E-14 |
| 22083_31 | 0.6444671 | 3.38E-14 |
| 23974_73 | 0.77487772 | 6.85E-16 |
| 26097_75 | 0.62594377 | 2.59E-11 |
| 26395_6 | 0.52172233 | 9.95E-10 |
| 27535_29 | 0.61752155 | 8.65E-13 |
| 27711_54 | 0.68995945 | 7.36E-17 |
| 27856_29 | 0.51409768 | 1.38E-09 |
| 28679_68 | 0.57049497 | 1.01E-12 |
| 29035_60 | 0.61357249 | 1.08E-12 |
| 30694_77 | 0.73893339 | 3.32E-15 |
| 30954_6 | 0.69956088 | 2.55E-13 |
| 31220_53 | 0.75990266 | 1.13E-14 |
| 32788_74 | 0.49512809 | 5.96E-11 |
| 33977_9 | 0.79851355 | 1.27E-17 |
| 34287_79 | 0.7482871 | 4.18E-13 |
| 35049_76 | 0.8298488 | 6.35E-16 |
| 35447_51 | 1.32235291 | 7.87E-18 |
| 35501_59 | 0.4664849 | 1.09E-09 |
| 36654_66 | 0.77361903 | 2.74E-15 |
| 36783_53 | 0.60915957 | 1.42E-12 |
| 38615_31 | 0.71350818 | 6.35E-16 |
| 38700_21 | 0.57489713 | 1.49E-10 |
| 38792_8 | 0.63461868 | 4.71E-12 |
| 39977_62 | 0.45706277 | 1.91E-08 |
| 40393_57 | 0.71170283 | 2.14E-15 |
| 41625_15 | 0.50702657 | 1.13E-10 |
| 42338_39 | 0.63696828 | 1.71E-11 |
| 42785_44 | 0.67320635 | 5.66E-12 |
| 43266_6 | 1.02357602 | 6.65E-34 |
| 45979_12 | 0.47839671 | 1.28E-11 |
| 45973_78 | 0.82528235 | 6.38E-21 |
| 50035_59 | 0.73802031 | 8.14E-12 |
| 52786_50 | 0.67276406 | 6.35E-16 |
| 54506_10 | 3.38705865 | 1.35E-70 |
| 55476_6 | 0.51849573 | 4.44E-10 |
| 55807_60 | 0.6886648 | 1.75E-14 |
| 62914_54 | 0.8323658 | 1.39E-15 |
| 62973_22 | 0.6854629 | 6.00E-13 |
